# Evolution and diversity of the TopoVI and TopoVI-like subunits with extensive divergence of the TOPOVIBL subunit

**DOI:** 10.1101/2022.05.08.491100

**Authors:** Julia Brinkmeier, Susana M. Coelho, Bernard de Massy, Henri-Marc Bourbon

## Abstract

Type II DNA topoisomerases regulate topology by double-stranded DNA cleavage and ligation. The TopoVI family of DNA topoisomerase, first identified and biochemically characterized in Archaea, represents, with TopoVIII and mini-A, the type IIB family. TopoVI has several intriguing features in terms of function and evolution. TopoVI has been identified in some eucaryotes, and a global view is lacking to understand its evolutionary pattern. In addition, in eucaryotes, the two TopoVI subunits (TopoVIA and TopoVIB) have duplicated and evolved to give rise to Spo11 and TopoVIBL, forming TopoVI-like (TopoVIL), a complex essential for generating DNA breaks that initiation homologous recombination during meiosis. TopoVIL is essential for sexual reproduction. How the TopoVI subunits have evolved to ensure this meiotic function is unclear. Here, we investigated the phylogenetic conservation of TopoVI and TopoVIL. We demonstrate that BIN4 and RHL1, potentially interacting with TopoVIB, have co-evolved with TopoVI. Based on model structures, this observation supports the hypothesis for a role of TopoVI in decatenation of replicated chromatids and predicts that in eucaryotes the TopoVI catalytic complex includes BIN4 and RHL1. For TopoVIL, the phylogenetic analysis of Spo11, which is highly conserved among Eukarya, highlighted a eukaryal-specific N-terminal domain that may be important for its regulation. Conversely, TopoVIBL was poorly conserved and rapidly evolving, giving rise to ATP hydrolysis-mutated or -truncated protein variants, or was undetected in some species. This remarkable plasticity of TopoVIBL provides important information for the activity and function of TopoVIL during meiosis.

## Introduction

DNA topoisomerases play an important role in resolving topological constraints and were first discovered in bacteria almost 50 years ago (Gellert et al., 1976; Wang, 1971). Ever since, DNA topoisomerases have been identified in many other organisms, such as archaea, eukaryotes and viruses (Chen et al., 2013; Forterre and Gadelle, 2009; Gadelle et al., 2014). They are ubiquitously expressed and play an essential role in various processes, such as replication, transcription, recombination, repair and chromatin remodeling (Chen *et al*., 2013; Vos et al., 2011; Wang, 2002). DNA topoisomerases have been extensively studied, and new family members have been progressively identified, adding insights into their structures and catalytic mechanisms.

Based on these features, DNA topoisomerases have been classified into two major types: type I and type II. Type I topoisomerases can transiently induce a break in one DNA strand, followed by passage of a single strand and re-ligation. Type II topoisomerases transiently break both DNA strands in a DNA double helix, followed by double-strand passage and break re-ligation (Champoux, 2001; Chen *et al*., 2013; Wang, 2002). Type II topoisomerases can relax negatively and positively supercoiled DNA, and have decatenation activity. Type II topoisomerases are further divided into type IIA and type IIB topoisomerases, and they both work in an ATP- and magnesium-dependent manner to resolve topological constraints during transcription, replication and recombination (Chen *et al*., 2013).

Topoisomerase VI (TopoVI) is a type IIB topoisomerase that was first identified in archaea (Bergerat et al., 1997) and forms a heterotetramer composed of two A and B subunits (*A*_2_*B*_2_) (Corbett et al., 2007; Graille et al., 2008). The A (TopoVIA) subunit carries the DNA cleavage activity and has two main domains: the 5Y-CAP (catabolite activator protein) DNA binding domain (also known as WHD, for winged helix domain), and the Toprim (topoisomerase-primase) domain (fig. 1A). The B (TopoVIB) subunit contains the GHKL domain with the ATP-binding/hydrolysis site, the helix-two-turn-helix (H2TH) domain, and the transducer domain that includes a long helical domain interacting with the N-terminal region of TopoVIA. The GHKL domain (ATPase) also known as the Bergerat fold (BF) contains three conserved motifs important for ATP binding that have been identified also in other proteins (i.e. DNA gyrase, heat-shock 90 family members, bacterial CheA histidine kinases, and MutL DNA mismatch protein family members) (Dutta and Inouye, 2000). Biochemical analyses of archaeal TopoVI showed that ATP binding is involved in TopoVIB homodimerization and plays an important role in activating the DNA cleavage complex. Three conserved basic residues within the so-called KGRR loop and C-ter Stalk/WKxY motif region of TopoVIB interact with DNA, with a preference for negative supercoiled DNA (Corbett *et al*., 2007; Wendorff and Berger, 2018). The KGRR loop is part of the GHKL domain, whereas the C-ter Stalk/ WKxY motif connects the transducer domain with TopoVIA and plays a role in DNA binding (Wendorff and Berger, 2018). Studies on the H2TH domain of TopoVI showed that it plays a role in facilitating DNA strand passage (Wendorff and Berger, 2018). The H2TH domain is followed by the transducer domain that contains a “switch lysine” required for ATP hydrolysis (Corbett et al., 2005).

**Figure 1.**
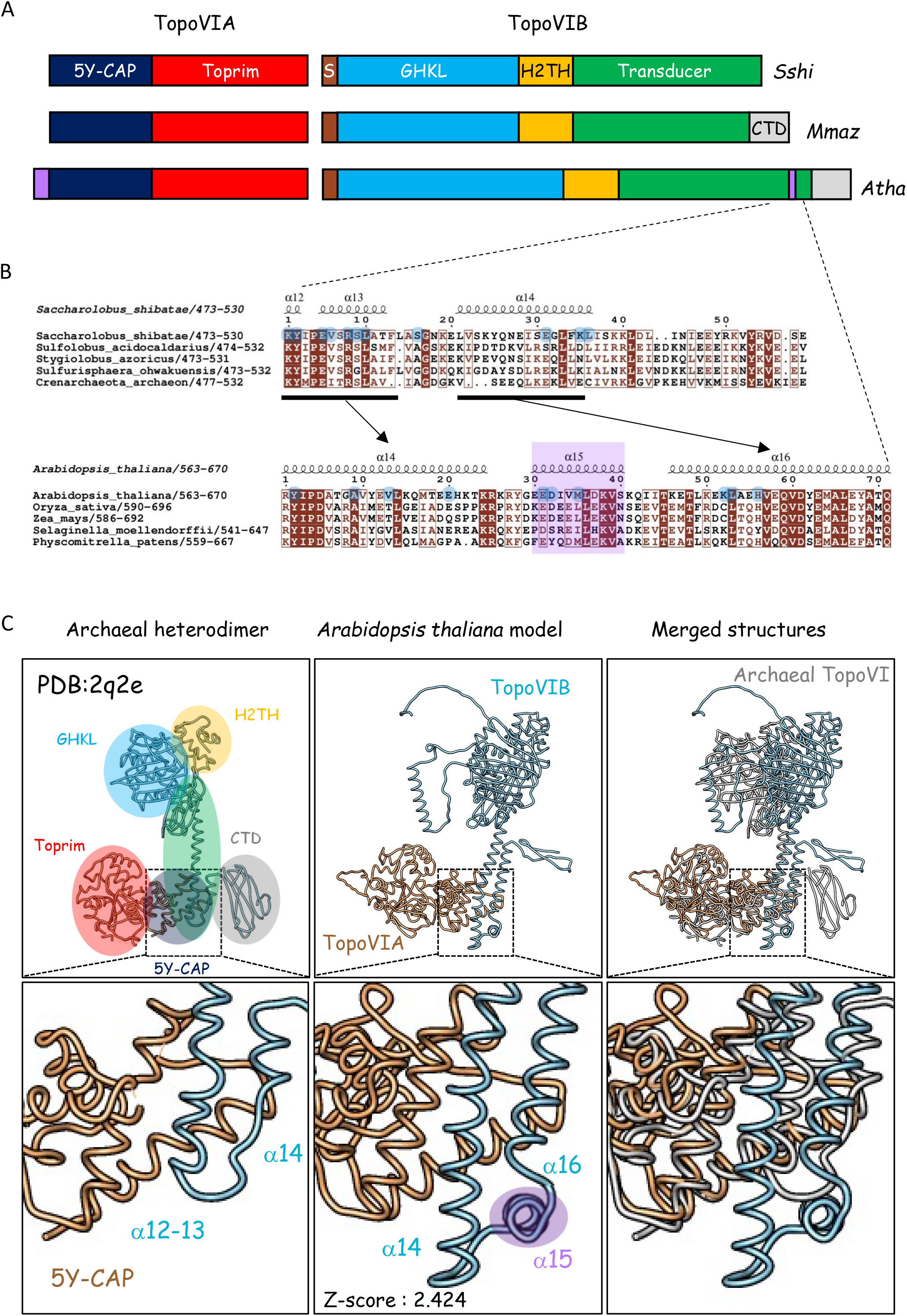
Archaeal and *A. thaliana* TopoVIB interact differently with TopoVIA. **A.** N- and C-terminal extensions in the eukaryotic TopoVIA and TopoVIB subunits. The domain organization of the *Saccharolobus shibatae (Sshi)*, *Methanosarcina mazei (Mmaz)*, and *Arabidopsis thaliana* (*Atha*) TopoVI subunits is shown. 5Y-CAP: catabolite activator protein, Toprim: topoisomerase-primase, S: N-terminal strap, GHKL: Gyrase B, Hsp90, Histidine Kinase, MutL, H2TH: helix-2turn-helix domain, CTD: C-terminal domain. The CTD is present in the TopoVIB of some archaea (*M. mazei*) but not in others (*S. shibatae*). The additional helices in *A. thaliana* TopoVIA and TopoVIB are highlighted in purple. **B.** Alignment of archaea and plantae TopoVIB transducer C-terminal subdomain showing the additional helix α15. Residues interacting with TopoVIA are highlighted with blue circles. *A. thaliana* TopoVIB structural elements are extracted from the AlphaFold database. **C.** Interaction of the TOPOVIBL transducer and TopoVIA 5Y-CAP in *M. mazei* (PDB:2q2e), and in *A. thaliana* (AF2 model). The whole structures are shown in the upper panels, with magnified views of the interacting helices in the lower panels. Archaeal TopoVIA/B domains are circled and coloured as in **A**.

In eukaryotes, TopoVI has been identified in vascular plants, green and red algae, and some protists (Hartung and Puchta, 2001; Malik et al., 2007b; Sugimoto-Shirasu et al., 2002; Yin et al., 2002). Differently from archaea, where TopoVI is the only Topo II activity (Forterre and Gadelle, 2009), eukaryotes also have Type IIA topoisomerases, raising the question of the functional specificity of TopoVI relative to Topo IIA. In plants, TopoVI is needed for DNA endoreduplication, potentially requiring a decatenation activity (Hartung et al., 2002; Kirik et al., 2007; Sugimoto-Shirasu *et al*., 2002; Yin *et al*., 2002). Potential TopoVI accessory factors (BIN4 and RHL1) have been identified in *Arabidopsis thaliana* mutants that have similar dwarf phenotypes (Breuer et al., 2007; Kirik *et al*., 2007; Sugimoto-Shirasu et al., 2005). Furthermore, various independent Yeast-two-Hybrid (Y2H) experiments indicate that BIN4 and RHL1 may be part of the TopoVI complex in *A. thaliana* (Breuer *et al*., 2007; Kirik *et al*., 2007; Sugimoto-Shirasu *et al*., 2005). The second specific feature of eukaryotes, compared with archaea, is the TopoVI-like (TopoVIL) complex that is composed of two proteins, SPO11 and TOPOVIBL, homologs to the TopoVIA and TopoVIB subunits (Bergerat *et al*., 1997; Keeney et al., 1997; Robert et al., 2016a; Robert et al., 2016b; Vrielynck et al., 2016). The TopoVIL complex induces DSBs at meiosis onset, an essential step to initiate homologous recombination and to create connections between homologous chromosomes and their proper segregation at meiosis I (Hunter, 2015). Although no biochemical information about its activity is currently available, *in vivo* data indicate that the TopoVIL complex can introduce DSBs, form a protein-DNA cleavage complex (like TopoVI), but differently from TopoVI, broken DNA ends are not predicted to re-ligated (Bergerat *et al*., 1997; Keeney *et al*., 1997). Indeed, the phospho-tyrosine linkage of the cleavage complex is not reversed, and SPO11 is released from substrate as a SPO11-oligo intermediate by endonucleolytic cleavage (Lange et al., 2011; Neale et al., 2005). The broken ends are repaired by homologous recombination (Baudat et al., 2013). It results that SPO11 is not recycled and undergoes only one catalytic rection. SPO11 is present in most eukaryotes examined and its phylogeny has been described (Bloomfield, 2016; Malik et al., 2007a; Ramesh et al., 2005), consistent with the meiotic pathway conservation for sexual reproduction. Conversely, TOPOVIBL conservation and phylogeny remain to be determined because so far, TOPOVIBL has been identified only in few species, partly due to the high degree of divergence observed among orthologs (Robert *et al*., 2016a).

Here, we extended the identification of TopoVI and TopoVIL families. For TopoVI, our analysis revealed new structural properties of TopoVIA and TopoVIB, and insights into the structural properties of BIN4 and RHL1 in eukaryotes. For TopoVIL, we revealed the widespread conservation of TOPOVIBL, with astonishing divergence, but a structurally conserved transducer subdomain that interacts with SPO11, particularly in metazoans. Lastly, we showed that REC114, a direct partner of TOPOVIBL, is broadly conserved in metazoans together with MEI4, a REC114 interacting protein.

## Results and Discussion

### Phylum- and species-specific conservation of TopoVIA, TopoVIB, BIN4 and RHL1 in eukaryotes

The phylogeny of TopoVI, the first identified family of type IIB topoisomerases, was first limited to Archaea and Viridiplantae (Forterre and Gadelle, 2009) and was then extended after the identification of other type IIB family members: TopoVIII, present in archaea and conjugative plasmids of bacteria (Gadelle *et al*., 2014), and miniA proteins often encoded by viruses (Takahashi et al., 2020).

The identified orthologs revealed the high conservation of both TopoVIA and TopoVIB subunits. Within the DNA binding domain (5Y-CAP or WHD) of the TopoVIA subunit, specific residues are conserved, particularly the catalytic tyrosine residue (Y). Within the Toprim domain, residues involved in Mg^++^ binding are conserved (DxDxxG). TopoVIA also includes a conserved helix at its N terminus involved in the interaction with TopoVIB. Moreover, it was proposed that a short conserved motif in the C-terminal Toprim subdomain, called T2BI-box (for Type IIB topoisomerases Interaction), is involved in its activity based on phylogenic studies of TopoVIA and miniTopoVIA variants (Takahashi *et al*., 2020). In TopoVIB, the conserved motifs include those in the GHKL domain for ATP binding and hydrolysis (BF motifs), and residues in the transducer domain that includes the interaction interface with TopoVIA. Several conserved motifs contribute to DNA binding, dimerization and ATP hydrolysis: the H2TH, the N-strap, a basic loop in the GHKL domain, the anchoring asparagine, the switch lysine, and the WKxY motif (Corbett and Berger, 2003; Wendorff and Berger, 2018).

Besides Viridiplantae, TopoVI homologs have been identified in several phyla among Eukarya (Bloomfield, 2016; Malik *et al*., 2007a). We re-investigated their phylogeny (figs. 1, 2 and Table 1) and detected two novel structural hallmarks of eukaryal TopoVI: i) an additional C-terminal helix (*A. thaliana* helix α15) in TopoVIB, located between the two helices of the transducer domain involved in the interaction with TopoVIA (fig. 1B), and ii) a helix (helix α1) in TopoVIA, located N-terminally to the helix involved in the interaction with TopoVIB (Supplementary fig. 1A). Based on a model structure of the *A. thaliana* TopoVIA-TopoVIB heterodimer, TopoVIB helix α15 leads to a partially overlapping but distinct interaction between the A and B subunits (fig. 1C). The function of the additional TopoVIA N-terminal helix (helix α1) is unknown. Based on model structures, it might interact with the Toprim domain adjacent to the T2BI box (Supplementary fig. 1B). Interestingly, these two helices we identified in Eukarya are also present in Asgard archaea, a sister-group of eukaryotes (Zaremba-Niedzwiedzka et al., 2017). Sequence alignments and model structures highlighted these similarities between Asgard and Eukarya (Supplementary figs. 2 and 3). This illustrates the common ancestry between Asgard and Eukarya and establishes that these two helices are not eukaryal-specific. Analysis of eukaryotes in which TopoVI is present (Archaeplastida, some opisthokonts and stramenopiles, Alveolata and Rhizaria (SAR)) showed that TopoVIB contained the motifs required for dimerization and ATP binding/hydrolysis (N-strap; and N, G1 and G2 boxes within the GHKL domain) (fig. 2), the H2TH domain, and a conserved transducer domain, including the anchoring asparagine, the WKxY motif, and the switch lysine (Supplementary fig. 4B). Similarly, in TopoVIA, the residues important for the catalytic activity, DNA interaction (notably Mg^++^ binding), and the T2BI box were well conserved from Archaeplastida to SAR (Supplementary fig. 4A). The eukaryal/Asgard-specific helices may represent interfaces for interaction with partners, thus potentially contributing to additional regulation of eucaryotic TopoVI compared with archaeal TopoVI. Apart from TopoVIA, no TopoVIB partner has been identified biochemically yet. However, two putative TopoVIA partners have been identified in *A. thaliana* based on genetic screens: BIN4 (also called MIDGET) and RHL1 (also called HYP7). Plants carrying mutations in either of these genes have the same phenotype as TopoVIA (SPO11-3) and TopoVIB mutants. Moreover, BIN4 interacts with itself, with RHL1 and with TopoVIA, while RHL1 also interacts with TopoVIA (Breuer *et al*., 2007; Kirik *et al*., 2007; Sugimoto-Shirasu *et al*., 2005). These observations suggested that the activity of plant TopoVI may require RHL1 and BIN4. To test this hypothesis, we analyzed RHL1 and BIN4 conservation in Eukarya and developed structural models.

**Figure 2.**
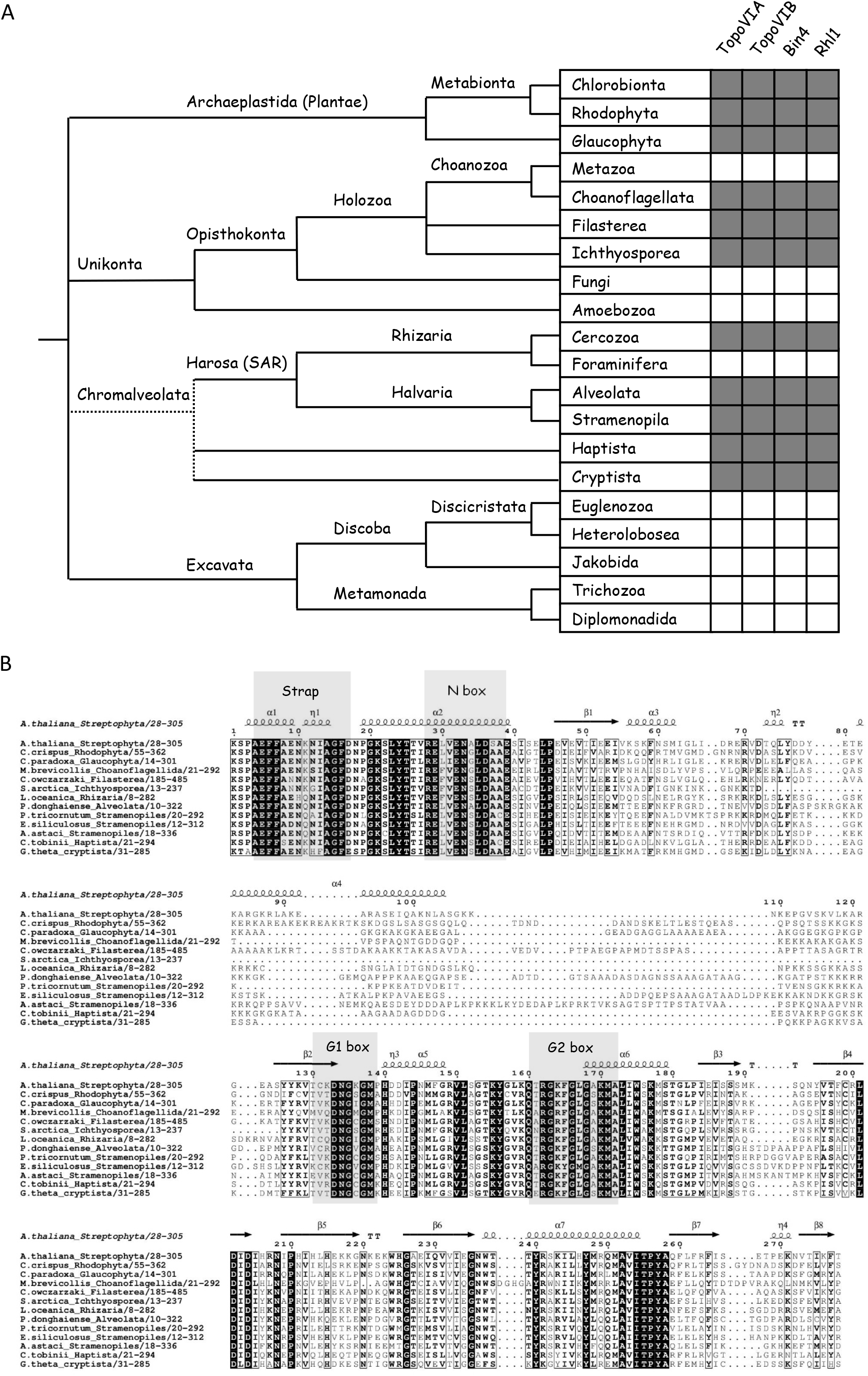
TopoVI holoenzyme subunit conservation among eukaryotic taxa. **A.** TopoVIA, TopoVIB, BIN4 and RHL1 conservation among Eukarya. The presence of TopoVIA, TopoVIB, BIN4 and RHL1 orthologs is shown by the gray boxes. The tree on the left side does not represent the genetic distances, because branch lengths are arbitrary and only used to illustrate the consensus view of the phylogenetic relationships among the major eukaryotic groups. Dashed lines indicate that chromalveolates are currently not considered to be monophyletic. **B.** Conservation of the BF motifs (N box, G1 and G2 boxes) in the GHKL domain of TopoVIB among Eukarya (Archaeplastida, Opisthokonta, SAR, Haptista and Cryptista). The alignment of the GHKL domain from eukaryal TopoVIB orthologs with the modeled structural elements from *A. thaliana* (extracted from the AlphaFold database) is shown. The N-strap (required for dimerization), and the N box and G boxes (required for ATP binding and hydrolysis) are highlighted in gray. Species: *Arabidopsis thaliana (Streptophyta)* *Chondrus crispus (Rhodophyta)* *Cyanophora paradoxa (Glaucophyta)* *Monosiga brevicollis (Choanoflagellata, Opisthokonta)* *Capsaspora owczarzaki (Filasterea, Opisthokonta)* *Sphaeroforma arctica (Ichthyosporea, Opisthokonta)* *Lotharella oceanica (Chlorarachniophyta, Rhizaria)* *Prorocentrum donghaiense (Dinophyta, Alveolata)* *Phaeodactylum tricornutum (Bacillariophyta, Stramenopila)* *Ectocarpus siliculosus (Phaeophyta, Stramenopila)* *Aphanomyces astaci (Oomycota, Stramenopila) Chrysochromulina tobinii (Haptista)* *Guillardia theta (Cryptista)*

**Figure 3.**
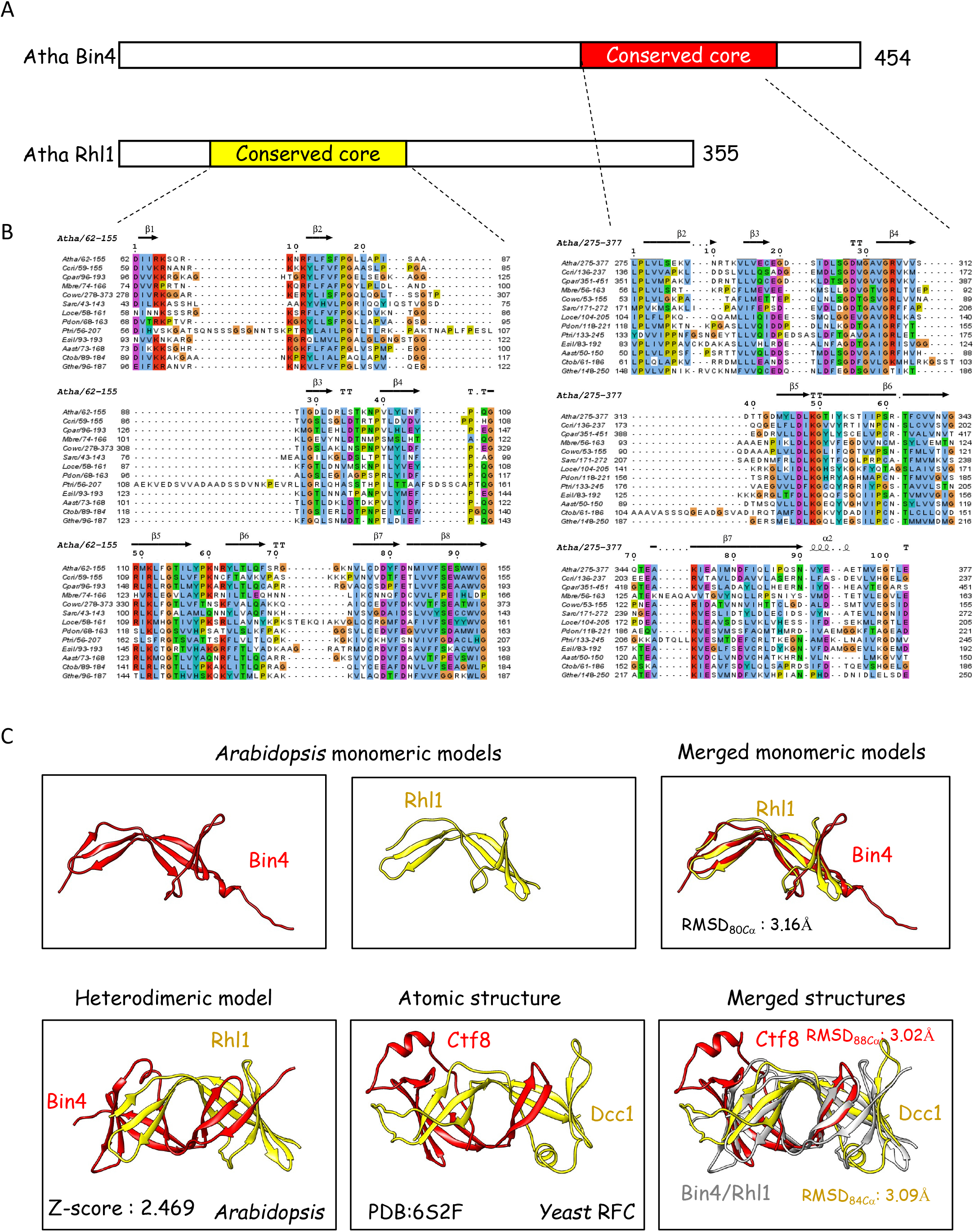
*A. thaliana* BIN4 and RHL1 interact through evolutionarily-conserved β-barrel domains that have structural similarities with the RFC Ctf8/Dcc1 dimerization module. **A.** Schematic representations of *A. thaliana* (Atha) BIN4 and RHL1, with the conserved cores highlighted in red and yellow, respectively. **B.** Alignment of the conserved cores of BIN4 and RHL1 among eukaryotes (same species as in Fig. 2), with the modeled structural elements from *A. thaliana* (extracted from the AlphaFold database). **C.** Structural AF2 models of *A. thaliana* BIN4 and RHL1 core domains. Upper panels: BIN4 (left) and RHL1 (middle) and merged (right) monomeric models. The docking root-mean-square-deviation (RMSD) value is indicated. Lower panels: Heterodimeric model of the BIN4/RHL1 dimerization module with the interaction Z-score (left). Atomic structure of the Ctf8/Dcc1 interface (PDB:6S2F) (middle). Merging of the Ctf8/Dcc1 and BIN4/RHL1 dimerization modules with the RMSD values (right).

**Table 1.** Summary of TopoVI and TopoVIL evolutionary conservation and diversity Domains and motifs present (+) or absent (-) or present in some phyla and absent in others (+/-) are shown for the two subunits of TopoVI (TopoVIA and TopoVIB) and of TopoVIL (Spo11 and TopoVIBL). na: non applicable due to lack of domain; L: linker domain substituting for the H2TH in TopoVIBL; 2H and 3H, 2 and 3 helices respectively.

|  |  | Nter helix | 5Y-CAP |  | TOPRIM |  | Cter |
| --- | --- | --- | --- | --- | --- | --- | --- |
|  |  |  | TopoVIB or BL-interacting domain | Catalytic Y | Mg++ binding | T2BI box |  |
| TopoVIA | Archaea | - | + | + | + | + | - |
|  | Asgard | + | + | + | + | + | - |
|  | Eukarya | + | + | + | + | + | - |
| Spo11 | Eukarya | +/- | + | + | + | + | *+ (*C. elegans*) |

|  |  | N strap | GHKL<br>Bergerat Fold (N, G1, G2 boxes) | H2TH | Transducer |  |  |  | CTD |
| --- | --- | --- | --- | --- | --- | --- | --- | --- | --- |
|  |  |  |  |  | Anchoring N | WKxY | Switch K | Spo11-interacting domain |  |
| TopoVIB | Archaea | + | + | + | + | + | + | + (2H) | +/- |
|  | Asgard | + | + | + | + | + | + | + (3H) | + |
|  | Eukarya | + | + | + | + | + | + | + (3H) | + |
| TopoVIBL | Eukarya | +/- | +/- | - (L) | +/- | + | +/- | + (3H) | + (Rec114 interacting domain) |
|  | GHKL | + | + | - (L) | + | + | + | + (3H) | + (Rec114 interacting domain) |
|  | GHKL-like | - | - | - (L) | - | + | - | + (3H) | + (Rec114 interacting domain) |
|  | No GHKL | na | na | - (L) | na | na | na | + (3H) | + (Rec114 interacting domain) |

This analysis led to three main conclusions: i) BIN4 and RHL1 are evolutionarily conserved in plants and also in some opisthokonts (including choanoflagellates, a sister group of metazoans) and in SAR. We did not find any BIN4 or RHL1 homolog in metazoans, fungi, amoebozoans, foraminiferans and excavates (fig. 2); ii) their presence is strictly correlated with that of TopoVI (A and B) (fig. 2); iii) analysis of BIN4 and RHL1 orthologs revealed for each protein a conserved central domain mostly composed of β-strands (fig. 3A, B). BIN4 and RHL1 models showed a striking structural overlap when monomers were compared (fig. 3C, upper right panel), as well as a dimeric structure (homo and hetero) in which the interaction between subunits involves the conserved domains (fig. 3C, lower left panel). BIN4 and RHL1 conserved domains shared similarity with the interacting β-barrel domains of the Ctf8/Dcc1 heterodimer (fig. 3C lower right panel). This heterodimer binds to the C-terminal end of Ctf18, a subunit of the RFC-CTF18 complex (Wade et al., 2017). Dcc1 also possesses three winged-helix domains (WHD), one of which binds ssDNA and dsDNA. The Ctf8/Dcc1 complex thus creates a bridge between DNA and RFC (Wade *et al*., 2017). The RFC-CTCF18 complex plays a role in loading PCNA at replication forks (Bermudez et al., 2003) and in sister chromatid cohesion (Kawasumi et al., 2021; Mayer et al., 2001). The similarity with BIN4 and RHL1 is thus particularly interesting since BIN4 and RHL1 also bind to DNA *in vitro* (Breuer *et al*., 2007; Sugimoto-Shirasu *et al*., 2005). Furthermore, we detected by modelling an interaction between the C-terminal domain of TopoVIB and the interface of β-strands from the BIN4/RHL1 heterodimer from *A. thaliana* (Supplementary fig. 5). To explore the evolutionary conservation of the interaction between BIN4 and RHL1, we used a Y2H assay and the model brown alga *Ectocarpus siliculosus* (Coelho et al., 2020). We found that *E. siliculosus* BIN4 and RHL1 interacted and that the conserved domain of each protein was necessary and sufficient for the interaction (Supplementary fig. 6). In *A. thaliana*, by Y2H assay, interactions were detected between TopoVIA and BIN4 or RHL1 (Breuer *et al*., 2007; Kirik *et al*., 2007; Sugimoto-Shirasu *et al*., 2005). We did not detect such interaction with *E. siliculosus* proteins, and based on our model (Supplementary fig. 5), we would predict an interaction with TopoVIB not TopoVIA.

The implication of these two putative TopoVIA partners could be important for the eukaryal TopoVI activity. As *A. thaliana* TopoVIA/B mutants are defective in endoreduplication, it was proposed that the TopoVI decatenation activity is involved in resolving catenated chromatids generated by endoreduplication. In fact, although TopoVI possesses both relaxation and decatenation activities, it was shown that *Methanosarcina mazei* TopoVI is preferentially a decatenase *in vitro* (McKie et al., 2022). Interestingly, the functional link with DNA replication is also supported by the interaction detected by immunoprecipitation between *Pyrococcus abyssi* TopoVI and the replication fork components PCNA, NusC and RFC (Ren et al., 2009). In Eukarya, some factors may also recruit TopoVI to replication forks, and BIN4 and RHL1 may stabilize such interactions.

### Early origin of the TopoVIL complex and TOPOVIBL divergence High conservation of SPO11 orthologs

SPO11 is conserved in Eukarya, in agreement with the observation that most eukaryotes undergo sexual reproduction (Malik *et al*., 2007b). Dictyostelids, which lack a *Spo11* gene, are the only known exception. This observation raises a yet unsolved puzzle for meiotic recombination, dependent on Spo11 in most species but not in dictyostelids (Bloomfield, 2018). We identified another clade where SPO11 is apparently absent: Strongyloididae, including *Strongyloides ratti* and related nematodes. It is thought that early after the emergence of the last eukaryotic common ancestor (LECA), its descendants possessed two SPO11 copies (SPO11-1 and SPO11-2) that are detected based on the conservation of the two major domains (5Y-CAP and Toprim). Loss of one copy occurred several times independently during evolution, and only some species have retained both paralogs (SPO11-1 and SPO11-2 in Plantae, and also in some amoebozoans, alveolates and excavates). Conversely, many eukaryotes have only one SPO11 (SPO11-1 in opisthokonts, some amoebozoans, rhizarians, haptophytes, cryptophytes and excavates; and only SPO11-2 in stramenopiles, red and green algae)(Malik *et al*., 2007b). SPO11 splice variants are conserved in plants (Ku et al., 2020; Sprink and Hartung, 2014), and in mammals (Romanienko and CameriniOtero, 1999). Interestingly these conserved splice variants correspond to two isoforms, one with and one without the N-terminal helix required for SPO11 interaction with TOPOVIBL. The variant without this helix should be catalytically inactive. In mice, expression kinetics data showed that the variant with the N-terminal helix (called SPO11β) is predominant at prophase onset, when meiotic DSBs form, and the variant without this helix (SPO11α) is predominant later in prophase (Bellani et al., 2010). Therefore, splicing regulation might contribute to modulate meiotic DSB activity.

In this new investigation of SPO11 orthologs, we identified SPO11 in all eukaryal supergroups (but for dictyostelids and possibly Strongyloididae), some with two paralogs, and with a high degree of conservation including the catalytic site (catalytic tyrosine, Mg^++^ binding site, and T2BI box) (figs. 4A, B, Table 1 and Supplementary fig. 7). Moreover, we found that a conserved additional N-terminal helix, absent in the characterized archaeal TopoVIA, and adjacent to the helix involved in TOPOVIBL interaction, was present in phyla-specific paralogs, reminiscent of the previously described additional N-terminal helix in Asgard archaea and eukaryal TopoVIA (Supplementary fig. 8A). This helix may bind to another partner and/or to the Toprim domain, as suggested by model structures for *A. thaliana* and Asgard TopoVIA (Supplementary figs. 1B and 2B). The only direct interacting partner of SPO11 identified (besides TOPOVIBL) is the *Saccharomyces cerevisiae* Ski8 protein that binds to the C-terminal domain of Spo11 (Arora et al., 2004). Interestingly, the Spo11 region interacting with Ski8 overlaps with the conserved T2BI motif (Takahashi *et al*., 2020) and is potentially interacting with the additional N-terminal helix of Spo11 (Supplementary fig. 1B). As some T2BI motif residues are next to the catalytic tyrosine in the *Methanococcus jannaschii* TopoVIA dimer in the absence of DNA, it was proposed that the T2BI motif is involved in regulating TopoVI activity (Takahashi *et al*., 2020). We infer that this is likely to apply also to TopoVIL.

We observed an exceptional feature of *C. elegans* SPO11: a sub-genus specific additional helix at the C-terminal end that might be implicated in partner interaction (Supplementary fig. 8). The presence of this extension in several *Caenorhabditis* species correlates with the apparent loss of TOPOVIBL. Indeed, structural modeling indicated that this extension is not present in three *Caenorhabditis* species that have TOPOVIBL homologs: *C. monodelphis*, *C. auriculariae*, and *C. parvicauda*.

### Rapid evolution and high diversity of TOPOVIBL homologs

TOPOVIBL phylogeny is more complex due to its high level of divergence, as already observed in a partial phylogenetic analysis within Eukarya (Robert *et al*., 2016a). In fact, we observed a large heterogeneity, from the presence of a conserved to a highly divergent B subunit to its non-detection. The divergence pattern suggests independent events, leading to the loss of similarity, and likely functional alterations relative to its TopoVIB ancestor subunit (figs. 4A, B, and Table 1). Importantly, it has been predicted that TopoVIL activity differs from that of TopoVI at least for the religation step, which is absent or repressed in the TopoVIL reaction (Robert *et al*., 2016b). Moreover, as some Type IIB topoisomerases can cleave DNA without ATP (Gadelle *et al*., 2014), the ATP binding and hydrolysis activity may not be an absolute prerequisite for TopoVIL function and/or this activity, which is also important for dimerization, may be substituted by alternatives molecular interactions.

**Figure 4.**
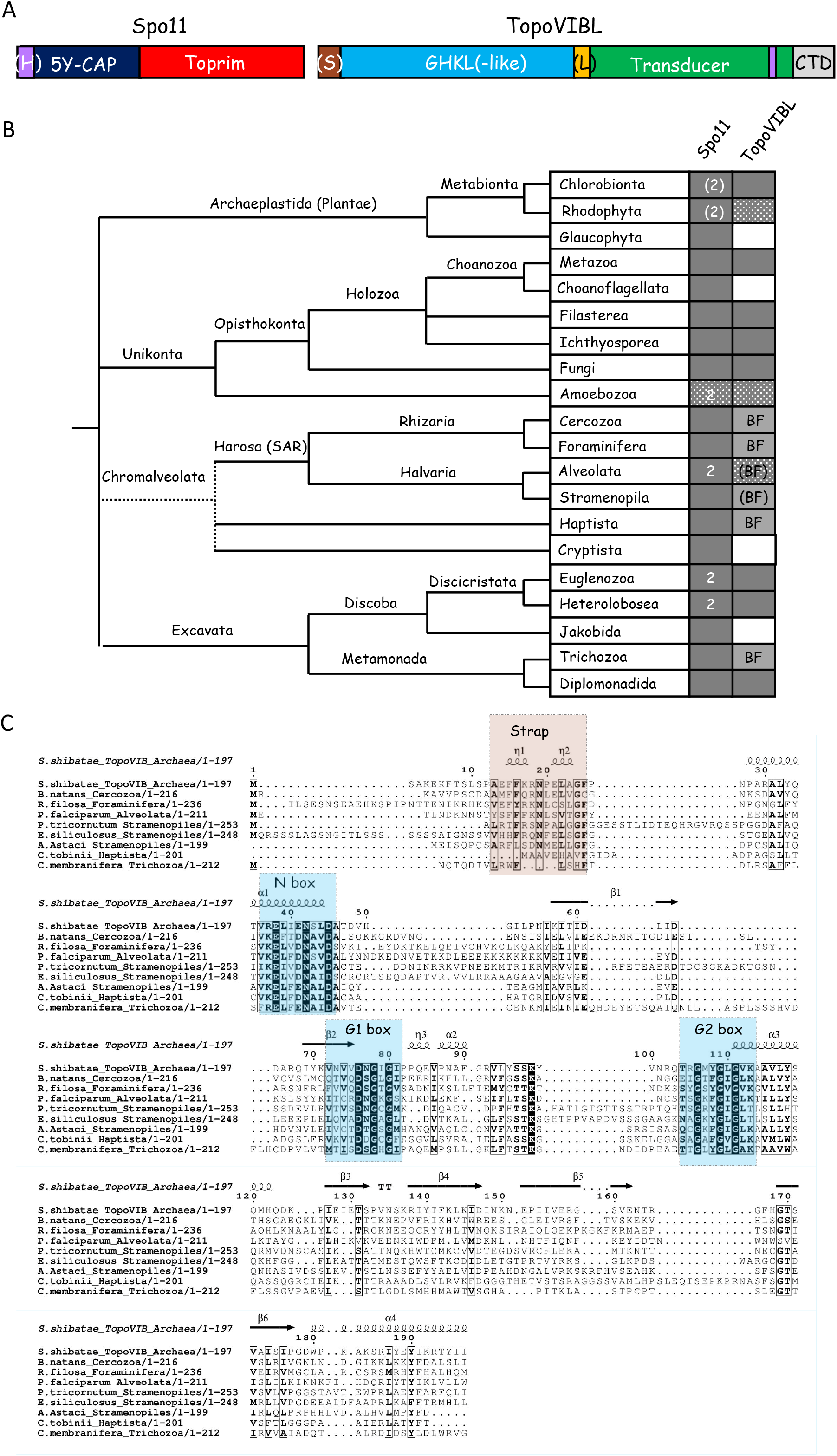
Phyla-specific conservation of the BF motifs of TOPOVIBL. **A.** Domains of SPO11 and TOPOVIBL. Some SPO11 orthologs have an additional N-terminal helix (H, purple box). The structure of TOPOVIBL orthologs is variable: with or without the N-Strap (S), GHKL domain with or without the conserved BF motifs (termed GHKL-like), and with a variable C-terminal domain (CTD). The region between the GHKL(-like) and the transducer domain is not conserved, and is called linker domain (L). **B.** Conservation of TopoVIL subunits among Eukarya. The presence (filled box) or partial occurrence (dotted box) of SPO11 and TOPOVIBL is indicated on the right. Some phyla have two SPO11 paralogues (2). In some phyla, TOPOVIBL contains conserved BF motifs. (BF) indicates BF motif heterogeneity among species within a phylum. A white box indicates that no TOPOVIBL ortholog could be identified. The phylogenetic relationships are described as in Fig. 2. **C.** Alignment showing the conservation of the N-strap and BF motifs in TOPOVIBL GHKL domains from SARs compared with archaeal TOPOVIB GHKL, with structural elements from *S. shibatae* (PDB: 2ZBK). Conserved boxes are highlighted as in Fig. 2. *Bigelowiella natans (Cercozoa, Rhizaria)* *Reticulomyxa filosa (Foraminifera, Rhizaria)* *Plasmodium falciparum (Apicomplexa, Alveolata)* *Phaeodactylum tricornutum (Bacillariophyta, Stramenopila)* *Ectocarpus siliculosus (Phaeophyta, Stramenopila)* *Aphanomyces astaci (Oomycota, Stramenopila)* *Chrysochromulina tobinii (Haptista)* *Carpediemonas membranifera (Trichozoa, Metamonada)*

In several phyla, TOPOVIBL is readily identifiable due to its high conservation with TopoVIB, specifically in the GHKL ATP binding/hydrolysis domain. The comparison of archaeal TopoVIB and TOPOVIBL from several chromalveolates highlighted the conservation of the N-strap required for dimerization, and of the N, G1 and G2 boxes required for ATP binding and hydrolysis (fig. 4C). In these species, the overall TOPOVIBL structure was highly similar to that of TopoVIB (Supplementary fig. 9), with the notable exception of the absence of the H2TH domain and the presence of an additional helix in the C-terminal transducer subdomain, compared with most archaeal TopoVIB. However, this additional helix was present also in TOPOVIBL and TopoVIB from Asgard archaea and eukaryotes (fig. 1 and Supplementary fig. 3). These highly conserved TOPOVIBL proteins can be differentiated from TopoVIB by the absence of the H2TH domain, which in TopoVI has DNA binding activity and is important for strand passage (Wendorff and Berger, 2018). Conversely, in other eukaryotic phyla, TOPOVIBL was highly divergent from TopoVIB. Specifically, they lost the N-strap, the three conserved boxes of the GHKL, the BF motifs (thus referred to as GHKL-like), and the H2TH domain. Therefore, the three helices within the C-terminal transducer subdomain that interacts with SPO11 were the major remaining conserved feature between these TOPOVIBL and TopoVIB (Supplementary fig. 10). The WKxY motif was only partially conserved, with mainly a remaining tryptophane residue (Supplementary fig. 10).

We observed the rapid evolution of TOPOVIBL with the specific loss of the conserved BF motifs in some phyla, such Oomycota (Stramenopila). Although some species, such as *Aphanomyces astaci* (Saprolegniales) had a highly conserved GHKL, in Albuginales (e.g., *Albugo candida* and *Albugo laibachii*), mutations disrupted both the N-strap and the conserved boxes (N, G1 and G2) required for ATP binding and hydrolysis. This indicates that TOPOVIBL function may have evolved by simultaneously losing a dimerization interface (the N-strap) and the ATPase activity (the BF motifs). However, the structural organization of the transducer region was conserved (Supplementary fig. 11). In *A. astaci*, in addition to the conservation of the GHKL BF motifs, models of the GHKL domain could be reconstituted with a high score when compared to *Saccharolobus shibatae* TopoVIB. This applied also to *A. astaci* TopoVIB and TOPOVIBL and showed specifically the conservation of the N-strap helix (Supplementary fig. 12). The ATP binding site also should be structurally conserved because its structural models were highly similar between *A. astaci* TopoVIB and TOPOVIBL (Supplementary fig. 13).

The TOPOVIBL organization plasticity was further revealed by one additional evolutionary outcome: the separation of the GHKL-like domain and the transducer in some species, for instance *S. cerevisiae*. In *S. cerevisiae*, REC102 was identified as homologous to the TOPOVIBL transducer domain (Robert *et al*., 2016a). It was then proposed that REC104, which does not contain any identifiable feature of a GHKL domain but which interacts with REC102, could substitute the GHKL domain (Robert *et al*., 2016b). This was shown by the biochemical characterization of the REC102/REC104 complex (Claeys Bouuaert et al., 2021b). Although we could detect truncated TOPOVIBL proteins, it is currently very challenging to determine in an unbiased manner whether an additional partner is present in a given species to replace the GHKL domain. REC104 could be identified as a partner, based on the genetic data that allowed showing its essential role for meiotic DSB formation in *S. cerevisiae* (Galbraith and Malone, 1992).

The high TOPOVIBL diversity was recapitulated in metazoans: few phyla have a TOPOVIBL with a GHKL domain, albeit lacking *bona fide* BF motifs, whereas many have a TOPOVIBL with a degenerated GHKL domain, named GHKL-like (with a conserved 3D fold, as determined by modeling), while retaining structural features in the transducer domain, specifically the three conserved helices (3H) predicted to be involved in the interaction with SPO11 (fig. 5). These metazoan TOPOVIBL proteins also contained a C-terminal domain (CTD) that includes an alpha helix interacting with REC114 in *M. musculus* TOPOVIBL (Nore et al., 2021) (Supplementary fig. 14). REC114 is an evolutionary conserved protein essential for meiotic DSB formation (Kumar et al., 2010). Its direct interaction with the TopoVIL complex in *S. cerevisiae* and *M. musculus* suggests that it plays a direct role in promoting DSB activity (Claeys Bouuaert et al., 2021a; Nore *et al*., 2021). An even more drastic evolutionary step is the complete loss of the GHKL-like domain, previously described in *S. cerevisiae*, *Schizosaccharomyces pombe* and *Drosophila melanogaster* (Robert *et al*., 2016a). We detected TOPOVIBL lacking any GHKL-like domain in several distant clades, such as Hemichordata, Platyhelmintha, Insecta, and Cnidaria (fig. 5B), and also within groups, such as in Amphibia (Chordata), Polychaeta (Annelida) and Chelicerata (Arthropoda)(Supplementary fig. 15). This indicates the very rapid evolution of this protein domain. Therefore, in these species, TOPOVIBL is only composed of a truncated transducer domain, limited to the 3H SPO11-interacting domain, and a C-terminal domain (fig. 5A). We identified a remarkably short TOPOVIBL form in *Schmidtea mediterranea*, where it is essentially composed of the 3H domain and a predicted C-terminal helix (fig. 5A and Supplementary fig. 15). Even in the absence of a GHKL-like domain, the interaction between the 3H domain and SPO11 could be modeled in *Pristionchus pacificus* and *C. monodelphis* (Supplementary fig. 16). As indicated in the Metazoa tree (fig. 5B), we could not detect any TOPOVIBL protein, even when searching for transducer-only proteins, in Placozoa (*Trichoplax adhaerens*) and the nematode *C. elegans*. These species may have lost not only the domain of interaction with SPO11, but also the one predicted to interact with REC114. TOPOVIBL remained undetected only in very close relatives to *C. elegans*, but not in the more diverging *C. monodelphis*, *C. auriculariae*, and *C. parvicauda*. In *C. elegans*, *C. auriculariae* and *C. parvicauda*, several meiotic genes, such as *dmc-1*, *mnd-1* and *hop-2*, have been lost (Rillo-Bohn et al., 2021)(our own observations); however, the loss of the gene encoding TOPOVIBL does not appear to be correlated. Understanding the potential implication of TOPOVIBL loss for SPO11 activity would require first to determine the precise role of the interactions between TOPOVIBL/SPO11 and TOPOVIBL/REC114. As biochemical information is available only for archaeal proteins, one could hypothesize that in eukaryotes these interactions may act to modulate SPO11 dimerization and/or binding to DNA, and/or the catalytic site position. In Placozoa and *C. elegans*, alternative strategies may have been selected, for instance by involving other partners for binding to SPO11. Interestingly, while the placozoan *T. adhaerens* has a single REC114 homolog, *C. elegans* has two REC114 paralogs (DSB-1 and DSB-2) of which one (DSB-1) interacts with SPO11-1 in Y2H assays (Hinman et al., 2021). This suggests that REC114 function is maintained, but acts by interacting with SPO11 rather than with TOPOVIBL.

**Figure 5.**
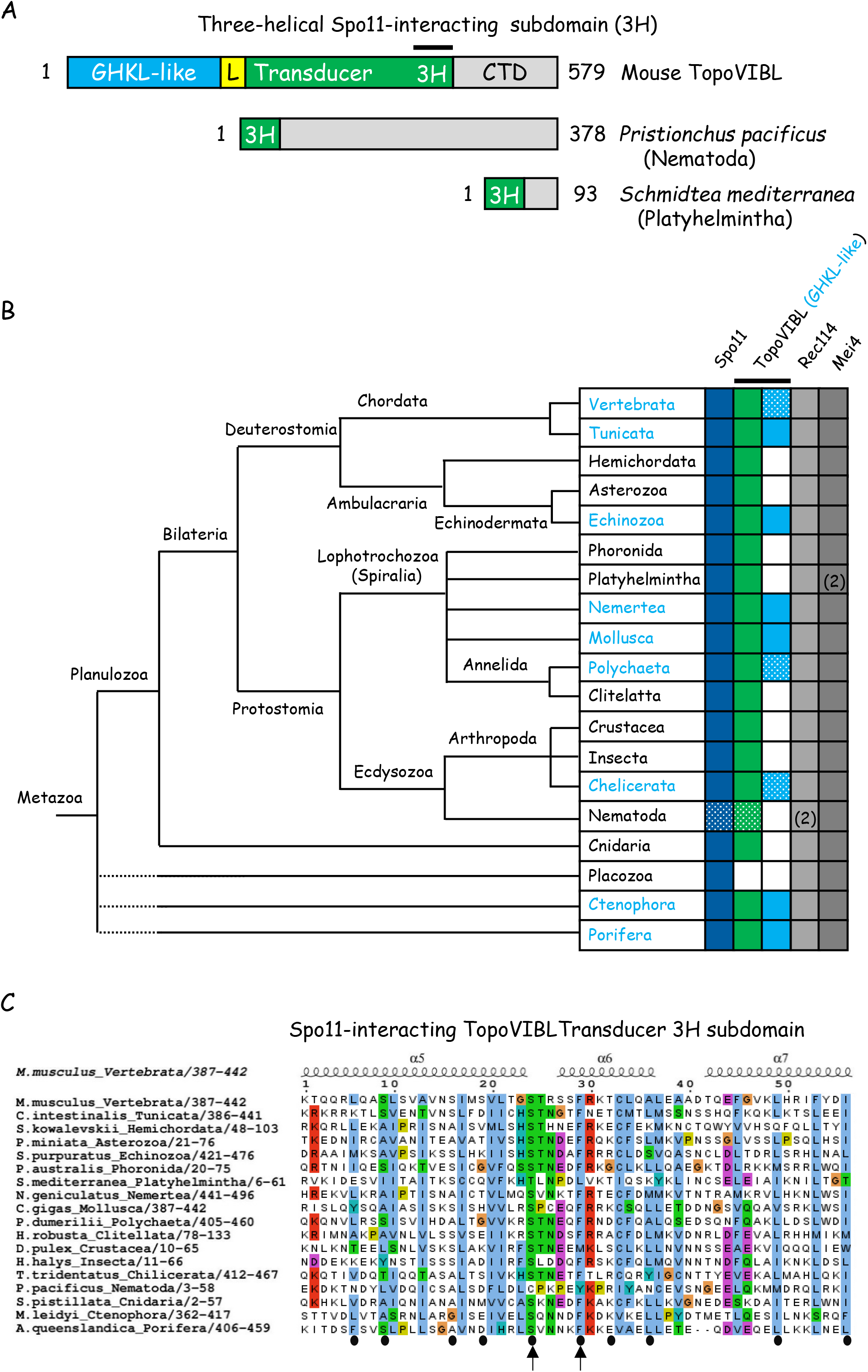
Conservation of meiotic DSB proteins among metazoans: phyla-specific loss of the TOPOVIBL GHKL-like domain and drastic reduction of the transducer domain. **A.** Schematic representation of metazoan TOPOVIBL proteins with a GHKL-like domain, a transducer including the three helices (3H) that interact with SPO11 and a C-terminal domain (CTD) (*M. musculus*), or with only the 3H and a CTD (*P. pacificus*), or with the 3H and a minimal CTD including one helix that might interact with REC114 (see alignment in Supplementary figs. S13-14) (*S. mediterranea*). **B.** Schematic phylogenetic tree showing the presence (filled boxes) or not (empty boxes) of SPO11, TOPOVIBL, REC114 and MEI4 in the indicated metazoans. Dotted boxes indicate heterogeneity (in the same phylum, some species contain and others do not SPO11 and/or TOPOVIBL). Phyla in which TOPOVIBL comprises a GHKL-like domain are indicated in blue letters and with blue filled boxes. Note that in some species among Vertebrata, Polychaeta and Chelicerata, TOPOVIBL orthologs lack the GHKL-like domain and have a transducer limited to the 3H domain (see Supplementary fig. S14). Phyla with species having two REC114 or MEI4 paralogs are indicated. The tree on the left side does not represent the genetic distances, because branch lengths are arbitrary and only used to illustrate the consensus view of the phylogenetic relationships among the major metazoan groups. Dashed lines indicate phyla for which the relationships are still debated. **C.** Conservation of the TOPOVIBL 3H domain. Alignment with structural elements of *M. musculus* predicted with AF2. The predicted residues interacting with SPO11 are indicated with black circles at the bottom of the alignment, and two strongly conserved residues (serine and phenylalanine) are indicated with black arrows. Species: *Mus musculus Ciona intestinalis* *Saccoglossus kowalevskii* *Patiria miniata* *Strongylocentrotus purpuratus* *Phoronis australia* *Schmidtea mediterranea* *Notospermus geniculatus* *Crassostrea gigas* *Platynereis dumerilii* *Helobdella robusta* *Daphnia pulex* *Halyomorpha halys* *Tachypleus tridentatus* *Stylophora pistillata* *Mnemiopsis leidyi* *Amphimedon queenslandica*

As mouse REC114 is a direct partner of TOPOVIBL (Nore *et al*., 2021) and of MEI4 (Kumar et al., 2018), we also explored its evolutionary conservation and identified REC114 and MEI4 orthologs in all metazoans examined (fig. 5, Supplementary figs. 17 and 18).

### Concluding remarks

The phylogenic reconstruction of TopoVI highlights a new and poorly explored property of TopoVI: the contribution of interacting partners. The concerted evolution of TopoVI with BIN4 and RHL1 suggests that these two proteins are directly involved in TopoVI catalytic activity. This information may help to understand why and how TopoVI has been retained in some eucaryotes. This could be a specific need linked to resolve catenated DNAs in DNA replication, in cells undergoing endoreduplication. Future functional and biochemical studies may answer these questions.

One TopoVI-specific, but not exclusive feature is its decatenase activity. As suggested by recent biochemical studies, this may be linked to specific substrate recognition. The proposition of TopoVI binding to DNA crossing is interesting with respect to the topological status of sister chromatids after DNA replication (McKie *et al*., 2022) and some factors may tether TopoVI to replication forks or intermediates. Understanding these TopoVI biochemical properties might also help to elucidate TopoVIL activity.

From *in vivo* studies in meiotic cells, it is clear that TopoVIL is distinct from TopoVI at least by the lack of or inactivated ability to religate DSBs.

TopoVIL activity takes place at meiotic prophase I onset, when DNA has been replicated. Although a temporal link with DNA replication has been shown in *S. cerevisiae* (Borde et al., 2000), the hypothesis that the TopoVIL substrate could be catenated sister chromatids is disfavored by the observation that in *S. cerevisiae* and *S. pombe*, DNA replication is dispensable (Blitzblau et al., 2012; Murakami and Nurse, 2001). TopoVIL activity is uncharacterized at the biochemical level, but distinct from TopoVI at least by the lack of or inactivated ability to religate DSBs. As it promotes DSBs, it is predicted to act as a dimer; however, the dimer stability should be required for break formation not for subsequent steps, unlike TopoVI. Thus, the contrast between the conservation of TopoVIB domains and the high diversity and evolution of TOPOVIBL is specifically striking (Table 1). The existence of TOPOVIBL orthologs with complete or partial loss of the GHKL domain seems to indicate that alternative molecular strategies have been developed for TopoVIL dimerization, and minimal versions of TopoVIBL seem to include a domain for interaction with Spo11, and a putative domain of interaction with Rec114. The contribution of partners such as Rec114, for regulating TopoVIL activity makes also sense since in the absence of religation activity, TopoVIL has to be turn on and off in a timely manner to avoid genome instabilities. The broad conservation of the associated proteins REC114 and MEI4 and of the REC114/TOPOVIBL interaction domains in Eukarya suggests that these proteins are directly involved in TopoVIL regulation.

## Materials and methods

### Alignments and models

Most TopoVIA, TopoVIB, BIN4, RHL1, SPO11, TOPOVIBL, REC114 and MEI4 homologs were identified from series of PSI-BLAST and HHMER analyses at the Max Planck Institute (MPI) using the MPI Toolkit (https://toolkit.tuebingen.mpg.de). Primary sequences were corrected through genomic translation and alternative exon prediction (https://web.expasy.org/translate/). Multiple sequence alignments (MSAs) generated by MAFFT 7.0 (http://mafft.cbrc.jp/alignment/server) with the auto mode and default parameters were used as inputs. MSA inputs included only previously validated homologs (as described hereafter), and alternative query sequences were systematically used as the MSA header sequence to improve detection of remote homologs. To discriminate among GHKL-containing homologs, MSA inputs included either full-length proteins or only their transducer domain, particularly the three-helical (H3) SPO11-interacting subdomain. Candidate proteins were validated or not by secondary structure prediction with PSI-PRED (http://bioinf.cs.ucl.ac.uk/psipred). Several DSB proteins were identified using PSI-BLAST or tblastn at NCBI (https://blast.ncbi.nlm.nih.gov/Blast.cgi), notably using transcriptome shotgun assembly (TSA) sequence databases, and dedicated BLAST servers. Remote homologs were validated by AlphaFold2 (AF2) modeling (see below), particularly on the basis of their capacity to stably interact with the SPO11 5Y-CAP domain through their H3 domain (Z-score>1.5). Protein-protein interaction Z-Scores were calculated with PIZSA (http://cospi.iiserpune.ac.in/pizsa/). MSAs were colored with JALVIEW 2.11.1.7 (https://www.jalview.org/) using the ClustalX similarity scheme, or with ESPript 3.0 (http://espript.ibcp.fr) using the %equivalent similarity score scheme, with the header sequence predicted secondary structure (as extracted from the AlphaFold database at the EBI website, or as determined from AF2 modeling, as described hereafter), and the Black&White (B&W) coloring scheme. Structural elements in figures 1B, 2B, 3A, 5C, S1, S2B, S4, S7-10, S14 and S17-18 were extracted from the AlphaFold database (https://alphafold.ebi.ac.uk/). Protein modeling was done with an advanced version of AF2 from ColabFold (https://github.com/sokrypton/ColabFold). Models are available upon request to HMB. Models were superimposed with MatchMaker from CHIMERA 1.15 (https://www.cgl.ucsf.edu/chimera/). ATP binding predictions were done with I-TASSER (https://zhanggroup.org/I-TASSER/). Protein structures were compared with DALI (http://ekhidna2.biocenter.helsinki.fi/dali/) and PDBeFold (https://www.ebi.ac.uk/msd-srv/ssm/).

### cDNA cloning

The potential *E. siliculosus* TOPOVI, BIN4 and RHL1 (protein and gene) sequences were identified based on a combination of sequence alignments (Henri-Marc Bourbon) and the RNA sequencing data available in the Genome viewer of OrcAE (*E. siliculosus* V2 data, https://bioinformatics.psb.ugent.be/orcae/). For the validation of the potential coding sequences of *E. siliculosus* genes, the cDNAs were synthesized by reverse transcription, as previously described in (Grey et al., 2016), using RNA samples from a diploid sporophyte (Ec702) and a haploid sporophyte (Ec32, male) undergoing apomeiosis (kindly provided by Susana Coelho, Station Biologique de Roscoff). This was followed by PCR amplification of the cDNA samples using gene-specific primers, as previously described in (Grey *et al*., 2016). The Gateway Gene Cloning system (Invitrogen) and synthesized cDNA (by GeneArt) optimized for expression in *E. coli* (compatible with codon usage of *S. cerevisiae*) were used for the cloning of full-length and truncated versions of *E. siliculosus* TOPOVI, BIN4 and RHL1 genes into pGADH-GW or pAS2dd.

### Yeast two hybrid assays

Yeast two hybrid assays were performed as previously described in (Imai et al., 2017). A diploid clone that expresses *M. musculus* Gal-4 AD-TOPOVIBL and Gal4 BD-SPO11β was used as a control for positive interactions (positive control) (Robert *et al*., 2016a).

## Acknowledgments

BdM was funded by ANR Topobreaks (ANR- 18-CE11-0024-01), Prize Coups d’Élan for French Research from the Fondation Bettencourt-Schueller, European Research Council (ERC) Executive Agency under the European Community’s Seventh Framework Programme (FP7/2007-2013 Grant Agreement no. [322788])), and CNRS. S. M. C was funded by ERC grant 864038. H-M. B was granted as a CNRS researcher. We thank Thomas Robert for discussions and critical reading of the manuscript.

## Supplementary figure legends

**Supplementary Figure 1 A.** Eukaryal TopoVIA has a conserved additional domain at its N-terminus. Alignment of Plantae and Archaea TopoVIA orthologs with modeled structure from *A. thaliana*. The additional domain (including the α1 helix) in Plantae is highlighted in purple. Three conserved motifs in the 5Y-CAP (in blue) and Toprim (in red) domains are highlighted: the catalytic site, the Mg^++^ binding site and the T2BI box.

AF2 modeling predicted potential interactions between the additional N-terminal domain and the Toprim domain through the residues highlighted as pink circles and as orange circles for each interaction site (see B). Red and pink arrows indicate interacting residues within the Toprim domain, next to the β-strand 13 and the T2BI box, respectively.

**B.** Model of the interaction within *A. thaliana* TopoVIA between the additional N-terminal domain and the Toprim domain, extracted from the AlphaFold database. Close-up view of the interaction next to β-strand 13 (left). *A. thaliana* TopoVIA model with the α1 helix (in purple) and unstructured neighboring residues (in yellow) (middle). Close-up view of the interaction next to the T2BI box (right).

**Supplementary Figure 2** Asgard TopoVIA also has an additional helical domain at its N-terminus.

**A.** Alignment of TopoVIA orthologs from *Lokiarchaeum sp. GC1475* and several uncultured *Candidatus Lokiarchaeota archaeon* species isolated from metagenomic samples, with the AF2 modeled structure from *Lokiarchaeum sp. GC1475*. The conserved catalytic site, Mg^++^ binding site, and T2BI box are highlighted with colored boxes as in Supplementary fig. S1.

**B.** Model of Asgard TopoVIA (left panel), and superposed to *A. thaliana* TopoVIA (right panel). The additional C-terminal helix is highlighted in purple, as depicted in A. Note that the Asgard and *A. thaliana* α1 helices are similarly modeled.

**Supplementary Figure 3** Asgard TopoVIB is predicted to interact with TopoVIA through a three-helix domain.

**A.** Alignment of the C-terminal region of TopoVIB orthologs from several *Candidatus Lokiarchaeota archaeon* metagenomic samples, with the AF2 modeled structure from the header sequence. The conserved WKxY motif (in green) and the additional helical domain (in purple) within the transducer C-terminal domain (CTD) (in gray) are highlighted. Note that the Asgard TopoVIA subunits have large CTDs with several structured motifs.

**B.** Models of the TopoVIB (blue)-TopoVIA(brown) interaction in Asgard (left panel) and in *A. thaliana* (central panel) and their superposition (right panel). The additional TopoVI alpha helix (in purple) is predicted to be part of a three-helix interaction domain with TopoVIA, both in Asgard and *A. thaliana*. Heterodimers were modeled with AF2.

.**Supplementary Figure 4** Conservation of the N-terminal helical extension (in purple) of TopoVIA (**A**), and of the H2TH (in yellow) and the additional helix (in green) within the transducer domain of TopoVIB (**B**) in Eukarya. Same species as in Fig. 2. The conserved catalytic site (in blue), Mg^++^ binding site, and T2BI box (both in red) of TopoVIA are highlighted. The WKxY motif is highlighted in green, and W and Y are the most conserved residues in TopoVIB orthologs. The conserved asparagine and lysine residues are indicated by arrows. Note that the TopoVIB CTD (in gray) is poorly conserved.

**Supplementary Figure 5.** Rhl1 and Bin4 are predicted to interact with the TopoVIB CTD and GHKL domains through their respective evolutionarily-conserved (Core) regions. Modelled interactions between *A. thaliana* and *E. siliculosus* TopoVI holoenzyme subunits (colored as in Figures 1 and 3), as determined by AF2. The predicted interaction between *A. thaliana* TopoVIB^CTD^ (circled in gray in the upper left panel) and Rhl1^Core^ is highlighted in gray (upper right panel). *A. thaliana* (left) and *E. siliculosus* (right) TopoVI heterodimeric complexes modelled with the Rhl1 (yellow) and Bin4 (red) Core domains are shown in the middle panels. The modelled interfaces between the TopoVIB CTD/GHKL domains and the Rhl1^Core^-Bin4^Core^ heterodimer are magnified in the lower panels, with the interacting regions highlighted in yellow and red ellipses, respectively.

**Supplementary Figure 6.** *E. siliculosus* BIN4 and RHL1 interact through their central conserved domains

**A.** Domains of TopoVIA, BIN4 and RHL1 from *E. siliculosus*

**B.** Summary of the Y2H interaction assay results. Five diploids (1 to 5) showed interaction between the proteins tested. FL: full length; +: growth on SDWLH-. Gray boxes: non-tested interactions.

**C.** Plate assays for the interactions tested by Y2H. Diploids 1 to 5, as in B. PC: positive control.

**D.** Western blots showing the expression of the indicated proteins used for the Y2H assays. Diploids 1 to 5, as in B. PC: positive control.

**Supplementary Figure 7.** SPO11 conservation among eukaryotic taxa. Alignment of SPO11 orthologs and paralogs with the modeled structure for *A. thaliana* SPO11.1, extracted from the AlphaFold database. The TOPOVIBL-interacting domain, the catalytic site (both in blue), the Mg^++^ interacting site and the T2BI box (both in red) are highlighted. Species: *Arabidopsis thaliana*

*Chondrus crispus*

*Cyanophora paradoxa*

*Monosiga brevicollis*

*Capsaspora owczarzaki*

*Amoebidium parasiticum*

*Saccharomyces cerevisiae*

*Planoprotostelium fungivorum*

*Bigelowiella natans*

*Reticulomyxa filosa*

*Plasmodium falciparum*

*Ectocarpus siliculosus*

*Chrysochromulina tobinii*

*Guillardia theta*

*Naegleria gruberi*

*Andalucia godoyi*

*Carpediemonas membranifera*

*Giardia intestinalis*

**Supplementary Figure 8.** *C. elegans* SPO11 displays a subgenus-specific C-terminal extension.

**A.** Alignment of SPO11 orthologs in metazoans with the modeled structure from the mouse header sequence, extracted from the AlphaFold database. Residues predicted to interact with SPO11 are highlighted with pink circles. The conserved additional N-terminal helix is highlighted in purple. The TOPOVIBL-interacting domain, the catalytic site (both in blue), the Mg^++^ interacting site, and the T2BI box (both in red) are highlighted as in fig. S6. The *C. elegans* additional C-terminal domain is highlighted in green. Species as in Fig. 5, except for *Eisenia fetida* (Clitelatta), *C. elegans* (Nematoda).

**B.** Alignment of the C-terminal part of SPO11 orthologs in different *Caenorhabditis* species, with the predicted modeled structure from the *C. elegans* header sequence, extracted from the AlphaFold database. Conservation is depicted with the ClustalX similarity coloring. Of note, the C-terminal extension is absent in the more diverging *C. monodelphis*, *C. auriculariae* and *C. parvicauda*. The conserved T2BI box is highlighted in red, and the predicted subgenus-specific C-terminal helix is highlighted in green.

**Supplementary Figure 9.** Conservation of the GHKL BF motifs in TopoVIB and TOPOVIBL among chromalveolates. Alignments of TopoVIB and TOPOVIBL paralogs with the modeled structure from the *A. thaliana* TopoVIB header sequence, extracted from the AlphaFold database, showing the conservation of the N-strap, N, G1 and G2 boxes (in blue), of the C-terminal additional helix and of the W residue within the WKxY motif (both in green). The positions of the conserved anchoring asparagine and switch lysine residues are indicated by vertical black arrows. Species:

*Bigelowiella natans (Cercozoa, Rhizaria)*

*Phaeodactylum tricornutum (Bacillariophyta, Stramenopila)*

*Ectocarpus siliculosus (Phaeophyta, Stramenopila)*

*Phytium insidiosum (Oomycota, Stramenopila)*

*Chrysochromulina tobinii (Haptista)*

**Supplementary Figure 10.** Overall TOPOVIBL conservation among Eukarya. Alignments of eukaryal TOPOVIBL orthologs with the modeled structure of the *M. musculus* header sequence, extracted from the AlphaFold database. The conserved GHKL(-like) domain is highlighted in blue. The WKxY motif (in green) can be identified in some species. The three-helix (3H) transducer subdomain predicted to interact with SPO11 is highlighted in green. Species:

*Arabidopsis thaliana* (Viridiplantae)

*Mus musculus* (Metazoa)

*Capsaspora owczarzaki (Filasterea)*

*Planoprotostelium fungivorum (Amoebozoa)*

*Bigelowiella natans (Rhizaria)*

*Reticulomyxa filosa (Foraminifera)*

*Plasmodium falciparum (Alveolata)*

*Phaeodactylum tricornutum (Stramenopila)*

*Ectocarpus siliculosus (Stramenopila)*

*Phytium insidiosum (Stramenopila)*

*Emiliania huxley (Haptista)*

*Leishmania infantum (Euglenozoa)*

*Naegleria gruberi (Heterolobosea)*

*Giardia intestinalis (Fornicata)*

**Supplementary Figure 11.** Rapid evolution of the TOPOVIBL GHKL domain: concomitant loss of the N-strap and BF motifs in Oomycota.

**A.** Alignments of oomycetal TOPOVIBL orthologs with the AF2 modeled structure from *A. astaci* TopoVIBL. The two Albuginales harboring degenerated N-strap and BF motifs are highlighted in pink.

**B.** Presence or absence of the TopoVI and TopoVIL subunits among oomycetes. This tree does not represent phylogenetic distances, because branch lengths are arbitrary and only used to separate diverse oomycetes. Colored boxes: presence of TopoVI/VIL subunits. Pink box: TOPOVIBL with degenerated BF motifs in albuginales.

**Supplementary Figure 12.** Oomycetal TopoVIB and TOPOVIBL are prone to dimerize through conserved N-straps. Upper panels, the structure of *S. shibatae* (left panel) TopoVIB GHKL (PDB:1MX0) is merged (right) with modeled *A. astaci* (middle) TopoVIB GHKL. Left panels, the structure of *S. shibatae* TopoVIB GHKL (PDB:1MX0) (upper) is merged (bottom) with the modeled *A. astaci* (middle) TOPOVIBL GHKL. Magnifications of the merged dimerized TopoVIB/VIBL interfaces are shown in the other panels, as indicated by dashed lines. Right bottom panel, the three magnified structures are merged showing the conservation of the helical N-straps, highlighted in brown. The interaction Z-scores are indicated.

**Supplementary Figure 13.** The *A. astaci* TopoVIB (upper panel) and TOPOVIBL (middle) GHKL domains are prone to ATP binding and hydrolysis as determined by structural modeling and molecular docking with AMP (I-Tasser models). Merged structures are shown in the lower panel.

**Supplementary Figure 14.** Weak conservation of the TOPOVIBL GHKL-like domain among metazoans. Alignment of TOPOVIBL orthologs with the modeled structure from *M. musculus* TOPOVIBL, extracted from the AlphaFold database. The GHKL-like domains (highlighted in blue) lack the most conserved residues of the N-Strap, N, G1 and G2 boxes. Conversely, the relatively-well conserved W residue within degenerated WKxY motif and SPO11-interacting domain (both in green) are present. The conserved REC114-interacting domain (in green) within the CTD (in gray) is highlighted. Note that AF2 modeling indicates properly folded GHKL domains, and that two (S and F) of the three (S, F and P) strictly conserved residues are detected within the SPO11-interacting transducer subdomain (see also Fig. 5C). Species:

*Mus musculus*

*Ciona savignyi*

*Strongylocentrotus purpuratus*

*Notospermus geniculatus*

*Crassostrea gigas*

*Platynereis dumerilii*

*Tachypleus tridentatus*

*Pleurobrachia bachei*

*Amphimedon queenslandica*

**Supplementary Figure 15.** Some metazoan TOPOVIBL orthologs lack the GHKL-like domain and have a truncated transducer domain that include only the SPO11-interacting 3H domain. **A.** Alignment of TOPOVIBL orthologs lacking the GHKL-like domain in different metazoan phyla, with the AF2 modeled structure from *Microcaecilia unicolor*, a primitive amphibian. The SPO11-interacting 3H domain and the C-terminal REC114-interacting domain are highlighted in green and gray, respectively. The alignment is colored using the ClustalX similarity scheme to emphasize structural conservation. Species:

*Microcaecilia unicolor*

*Saccoglossus kowalevskii*

*Patiria miniata*

*Phoronis australis*

*Schmidtea mediterranea*

*Lamellibrachia satsuma*

*Helobdella robusta*

*Thrips palmi*

*Ixodes scapularis*

*Caenorhabditis monodelphis*

*Euphyllia ancora*

**B.** Schematic phylogenetic tree showing the presence or absence (blue or white boxes, respectively) of the GHKL-like domain in TOPOVIBL (green filled boxes) among Amphibia, Polychaeta and Chelicerata (see Fig. 5B). This tree does not represent the phylogenetic distances, because branch lengths are arbitrary and only used to discriminate among clades.

**Supplementary Figure 16.** Modeled structures of TOPOVIBL-SPO11 interaction in *P. pacificus*, *M. musculus*, and *C. monodelphis*. AF2 models only include the TOPOVIBL 3H domain, and predict tight interaction with the helices α2-3 of SPO11. The interaction Z-scores and RMSD values (for the merged heterodimer structures) are indicated.

**Supplementary Figure 17.** REC114 conservation across Metazoa. Alignment of metazoan REC114 orthologs, with the modeled structure from *M. musculus*, extracted from the AlphaFold database. The conserved PH domain and the predicted MEI4-interacting domain are highlighted in green. Species:

*Mus musculus*

*Ciona intestinalis*

*Saccoglossus kowalevskii*

*Patiria miniata*

*Strongylocentrotus purpuratus*

*Phoronis australia*

*Schmidtea mediterranea*

*Notospermus geniculatus*

*Crassostrea gigas*

*Owenia fusiformis*

*Helobdella robusta*

*Daphnia pulex*

*Zootermopsis nevadensis*

*Parasteatoda tepidariorum*

*Pristionchus pacificus*

*Caenorhabditis elegans*

*Stylophora pistillata*

*Trichoplax adhaerens*

*Mnemiopsis leidyi*

*Amphimedon queenslandica*

**Supplementary Figure 18.** MEI4 conservation among Metazoa. Alignment of metazoan MEI4 orthologs, with the modeled structure from *M. musculus*, extracted from the AlphaFold database. The conserved predicted REC114-interacting domain is highlighted in green. Species as in Fig. S17.

