## Supplemental figures 1-18 for "Evolution and diversity of the TopoVI and TopoVI-like subunits with extensive divergence of the TOPOVIBL subunit"

### Figure S1

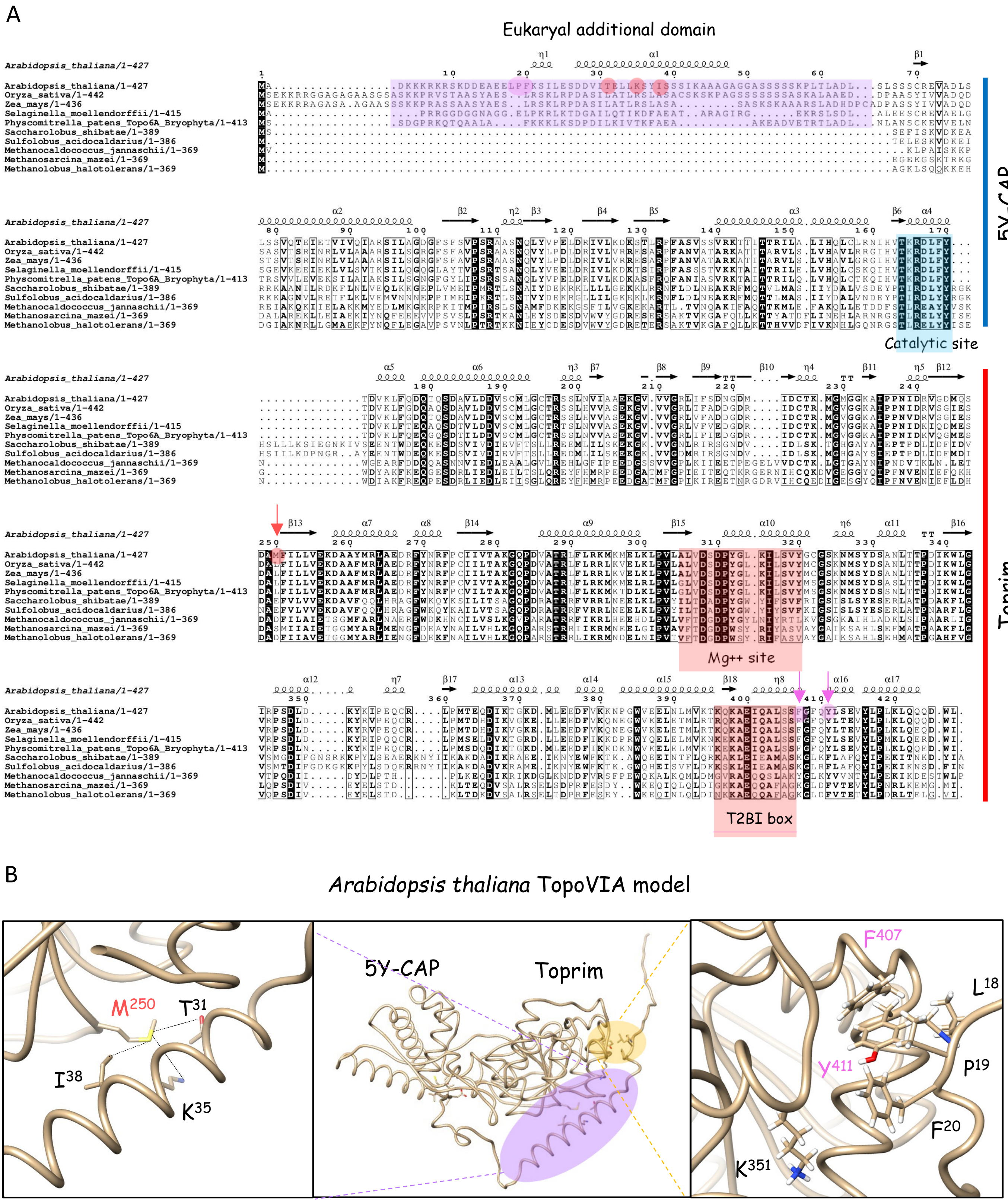
A
Eukaryal additional domain
Arabidopsis_thaliana/1-427
Oryza_sativa/1-442
Zea_mays/1-436
Selaginella_moellendorffii/1-415
Physcomitrella_patens_Topo6A_Bryophyta/1-413
Saccharolobus_shibatae/1-389
Sulfolobus_acidocaldarius/1-386
Methanocaldococcus_jannaschii/1-369
Methanosarcina_mazei/1-369
Methanolobus_halotolerans/1-369
5Y-CAP
Catalytic site
Toprim
Mg++ site
T2BI box
B
Arabidopsis thaliana TopoVIA model
M250
T31
I38
K35
5Y-CAP
Toprim
F407
L18
P19
Y411
F20
K351

### Figure S2

A

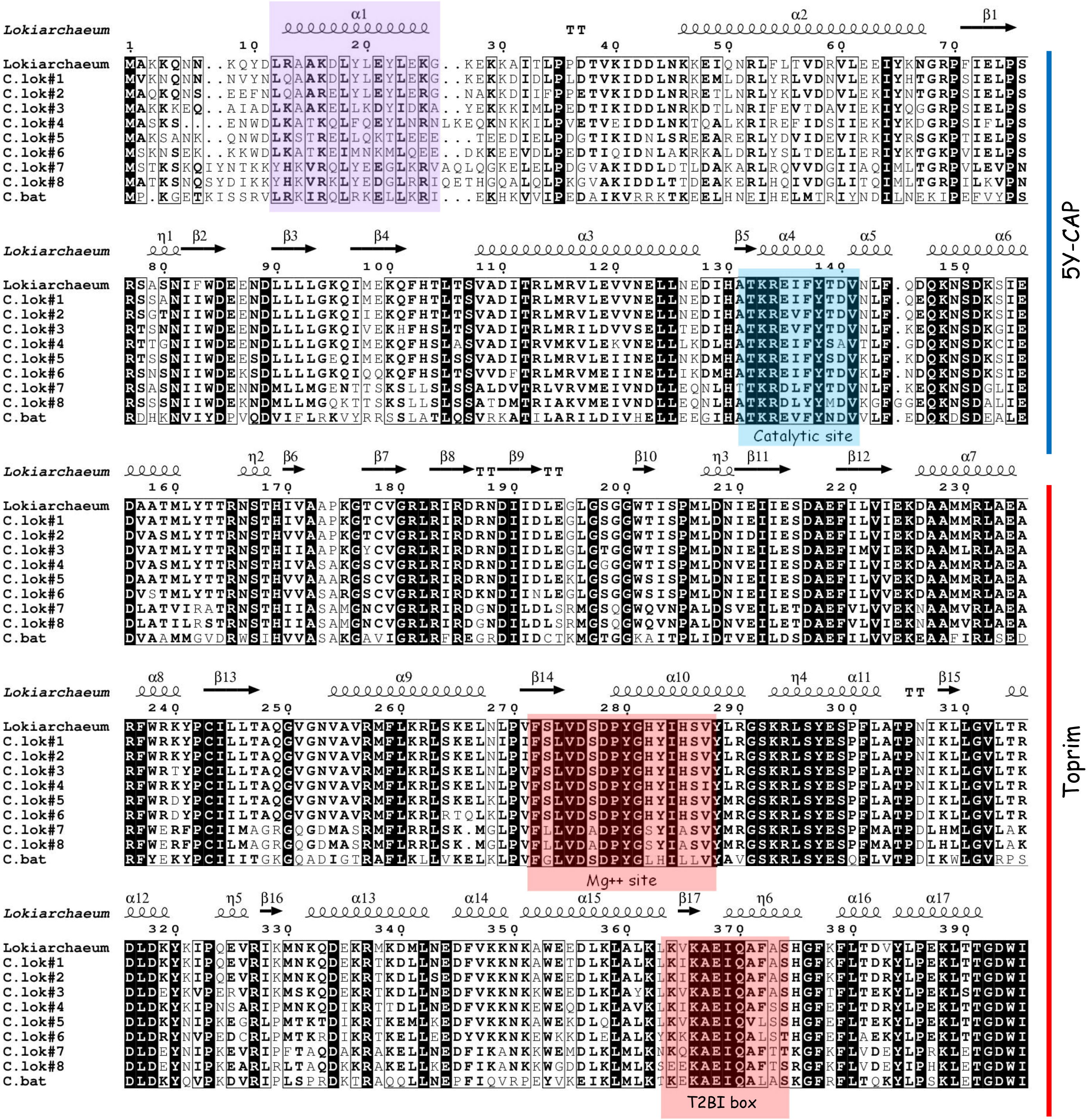


B

Asgard vs *Arabidopsis thaliana* TopoVIA models

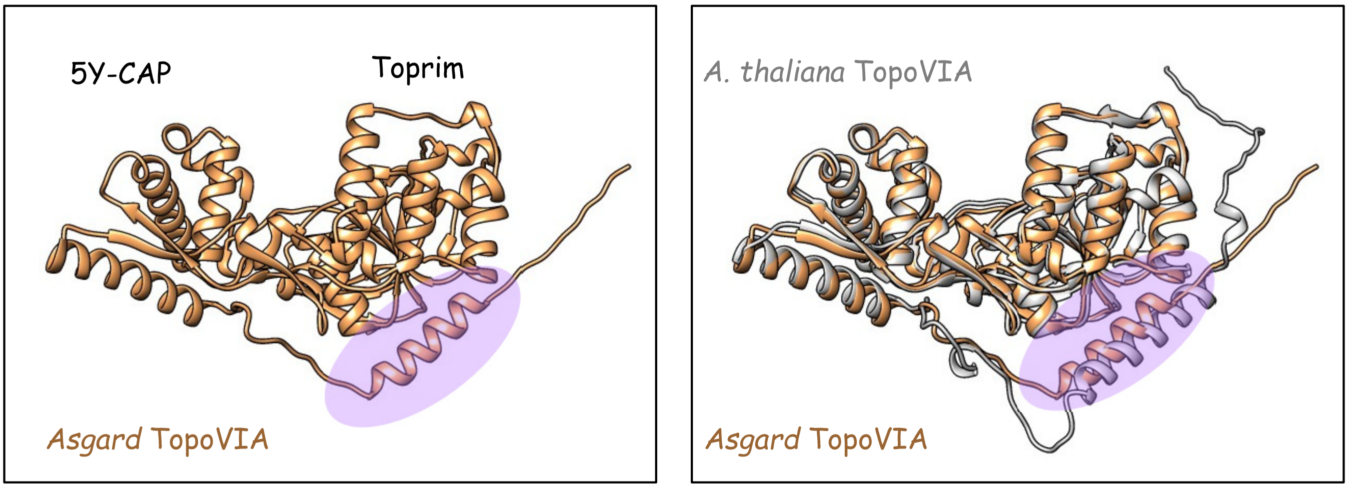

Figure S3

A

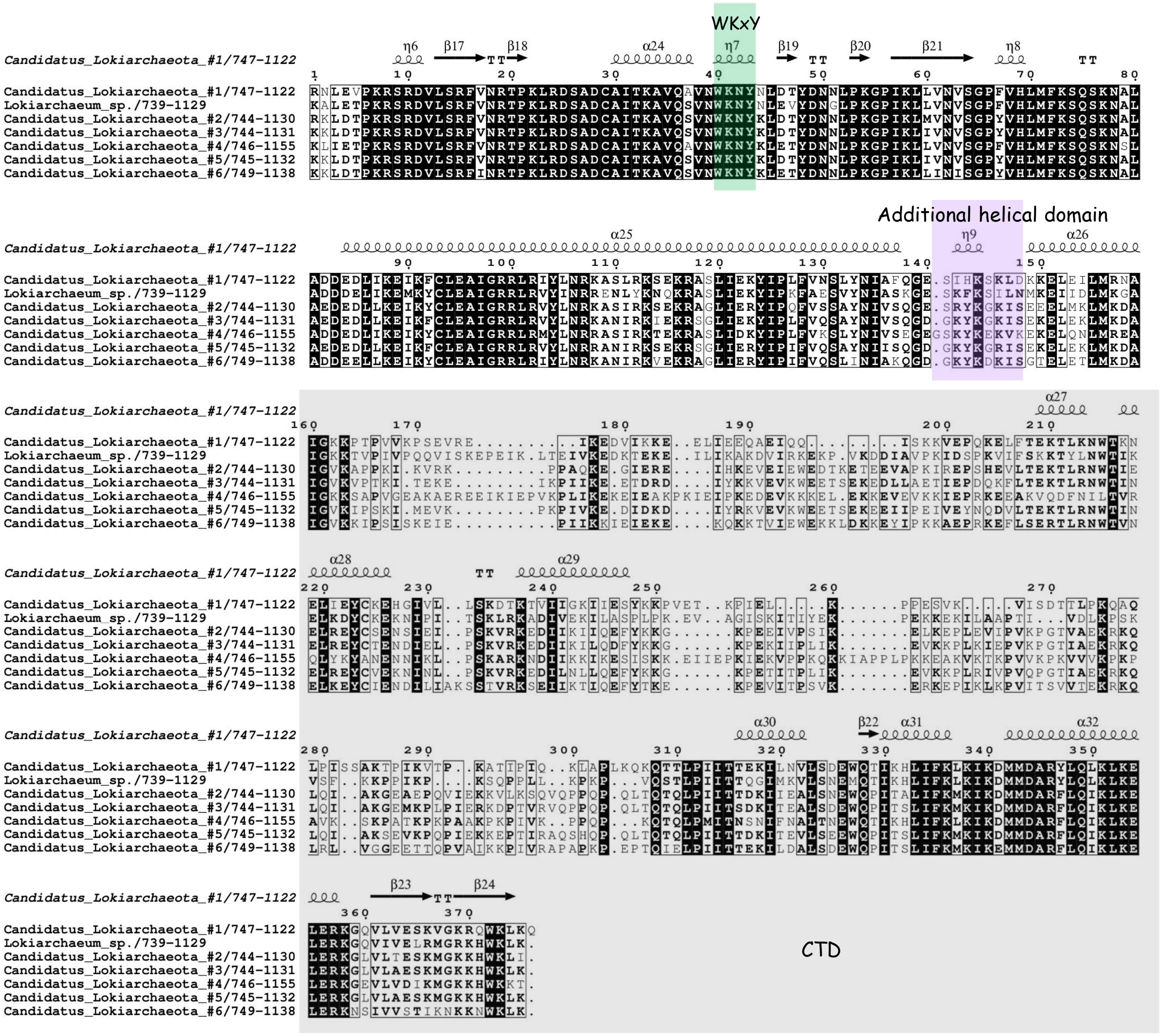


B

*Asgard vs Arabidopsis thaliana* TopoVIA/B heterodimer models

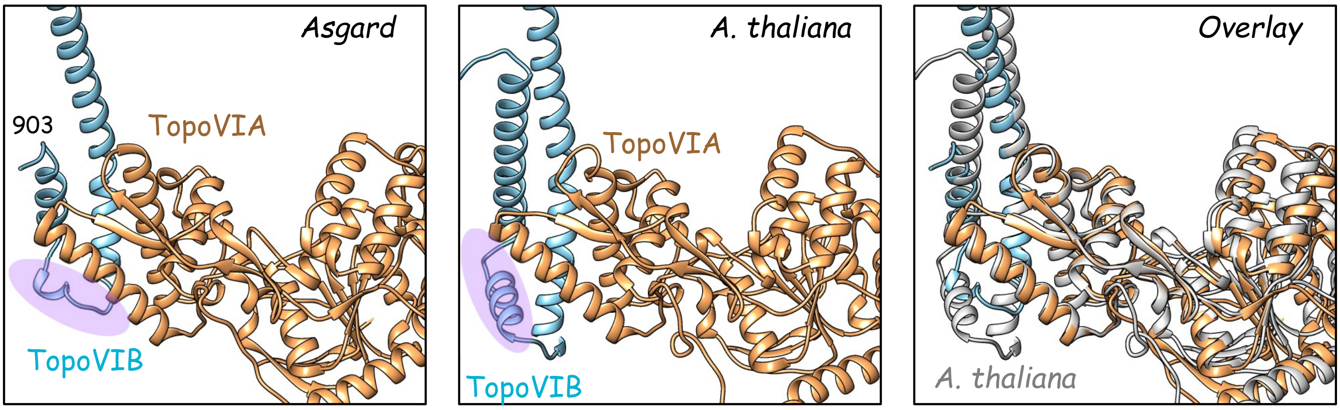

Figure S4

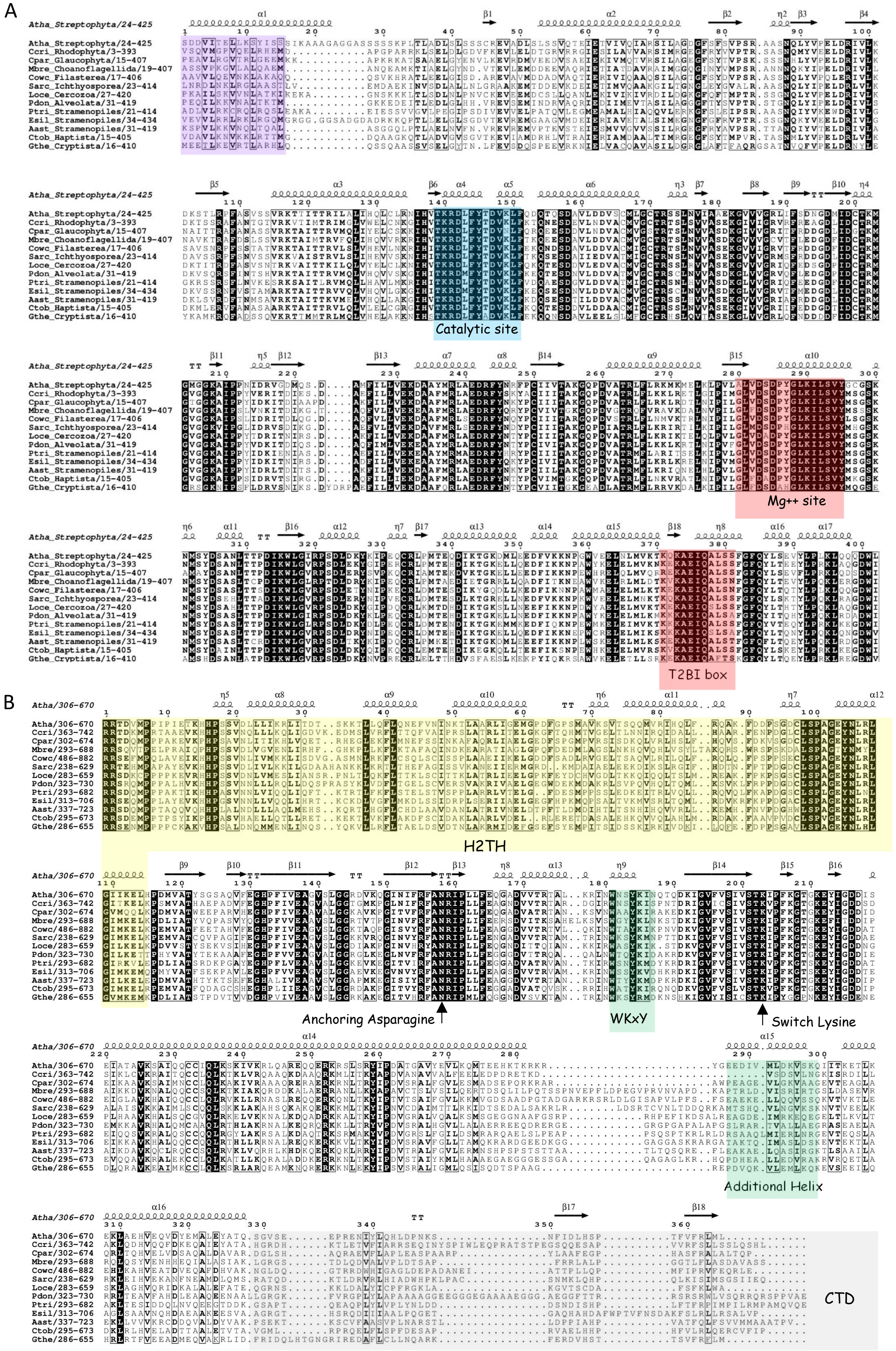
A
Catalytic site
Mg++ site
T2BI box
B
H2TH
Anchoring Asparagine
WKxY
Switch Lysine
Additional Helix
CTD

Figure S5

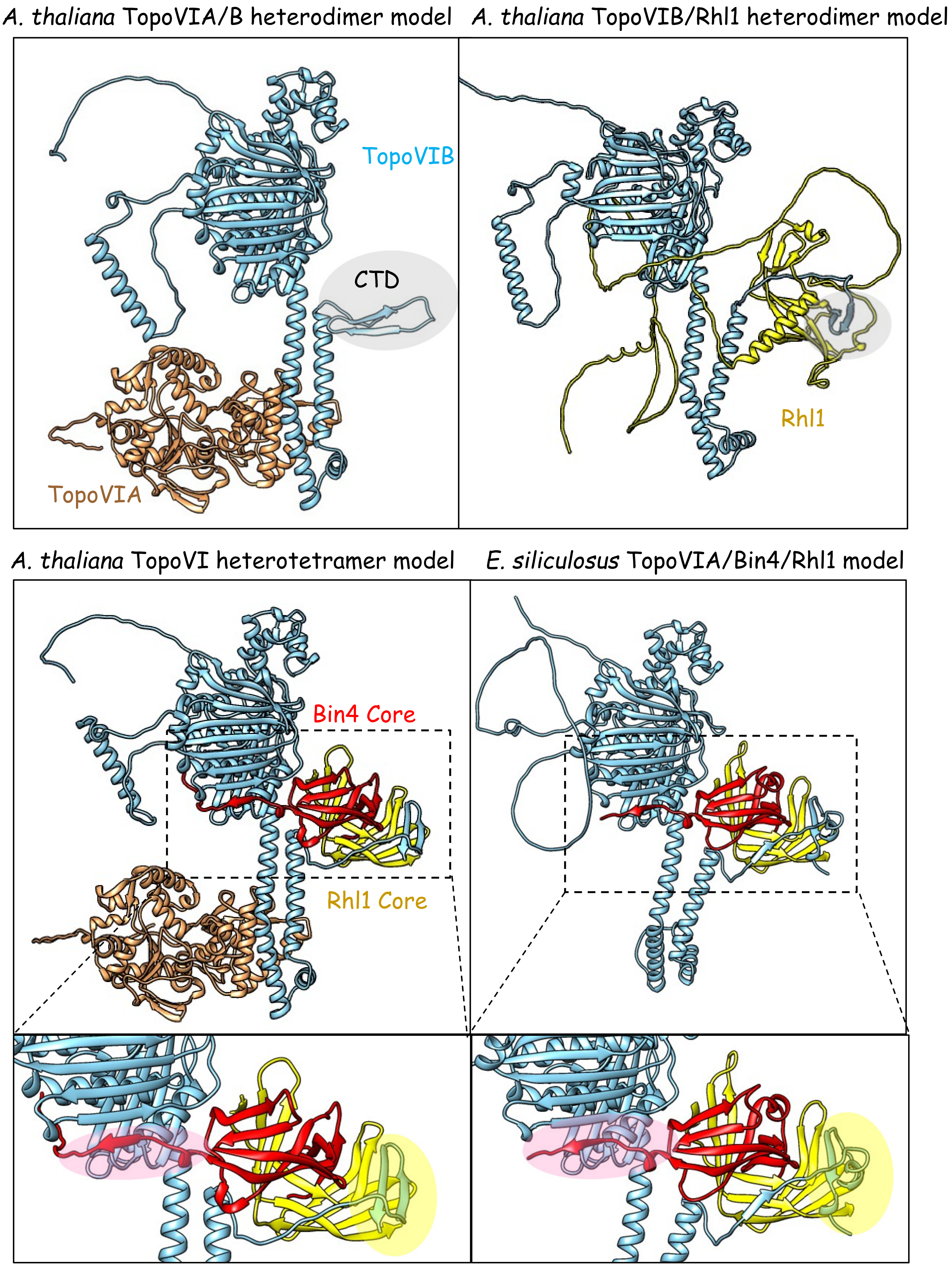

A. thaliana TopoVIA/B heterodimer model
A. thaliana TopoVIB/Rhl1 heterodimer model
TopoVIB
CTD
TopoVIA
Rhl1
A. thaliana TopoVI heterotetramer model
E. siliculosus TopoVIA/Bin4/Rhl1 model
Bin4 Core
Rhl1 Core

### Figure S6

A

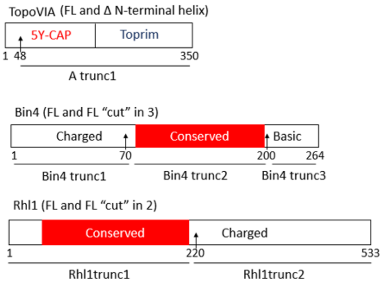


B

| pAS / pGAD | Bin4 FL | Bin4 trunc1 | Bin4 trun2 | Bin4 trunc3 | Rhl1 FL | Rhl1 trunc1 | Rhl1 trunc2 |
|---|---|---|---|---|---|---|---|
| TopoVIA FL | - | - | - | | - | | |
| TopoVIA trunc1 | - | - | - | - | - | - | - |
| Bin4 FL | - | | | | +[1] | +[2] | - |
| Bin4 trunc1 | | | | | - | - | |
| Bin4 trunc2 | | | | | - | +[4] | - |
| Bin4 trunc3 | - | | | - | - | - | - |
| Rhl1 FL | - | - | - | - | - | | |
| Rhl1 trunc1 | +[5] | - | +[3] | - | | | |
| Rhl1 trunc2 | | | - | - | | | |

C

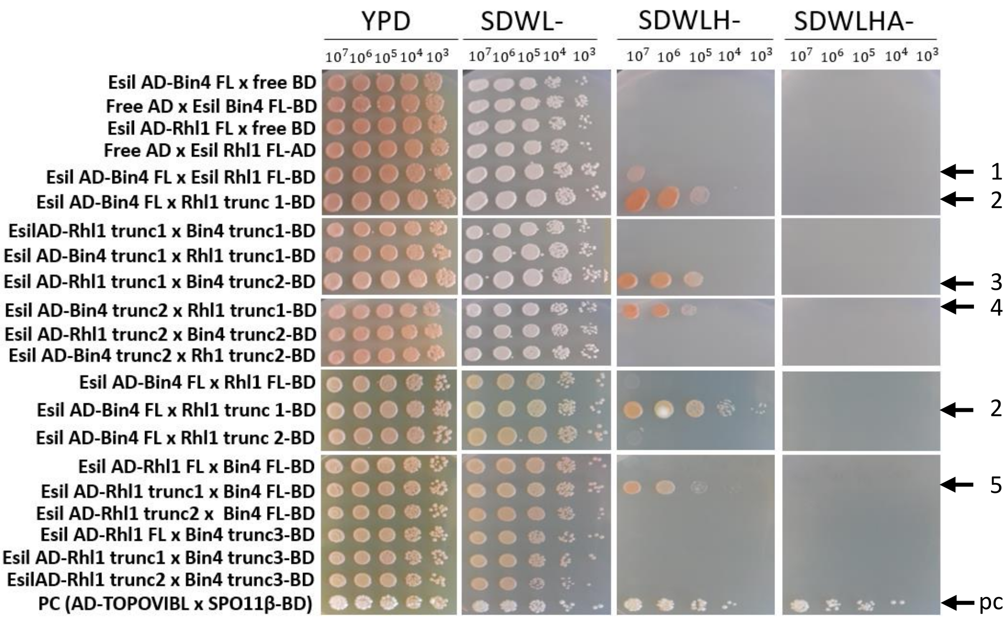


D

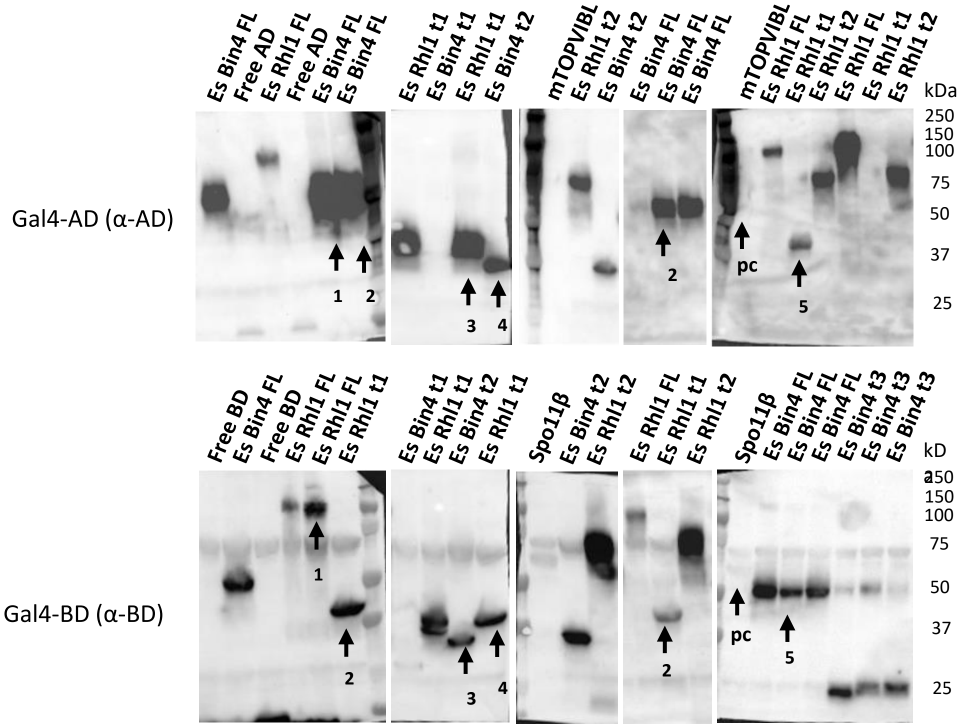

### Figure S7

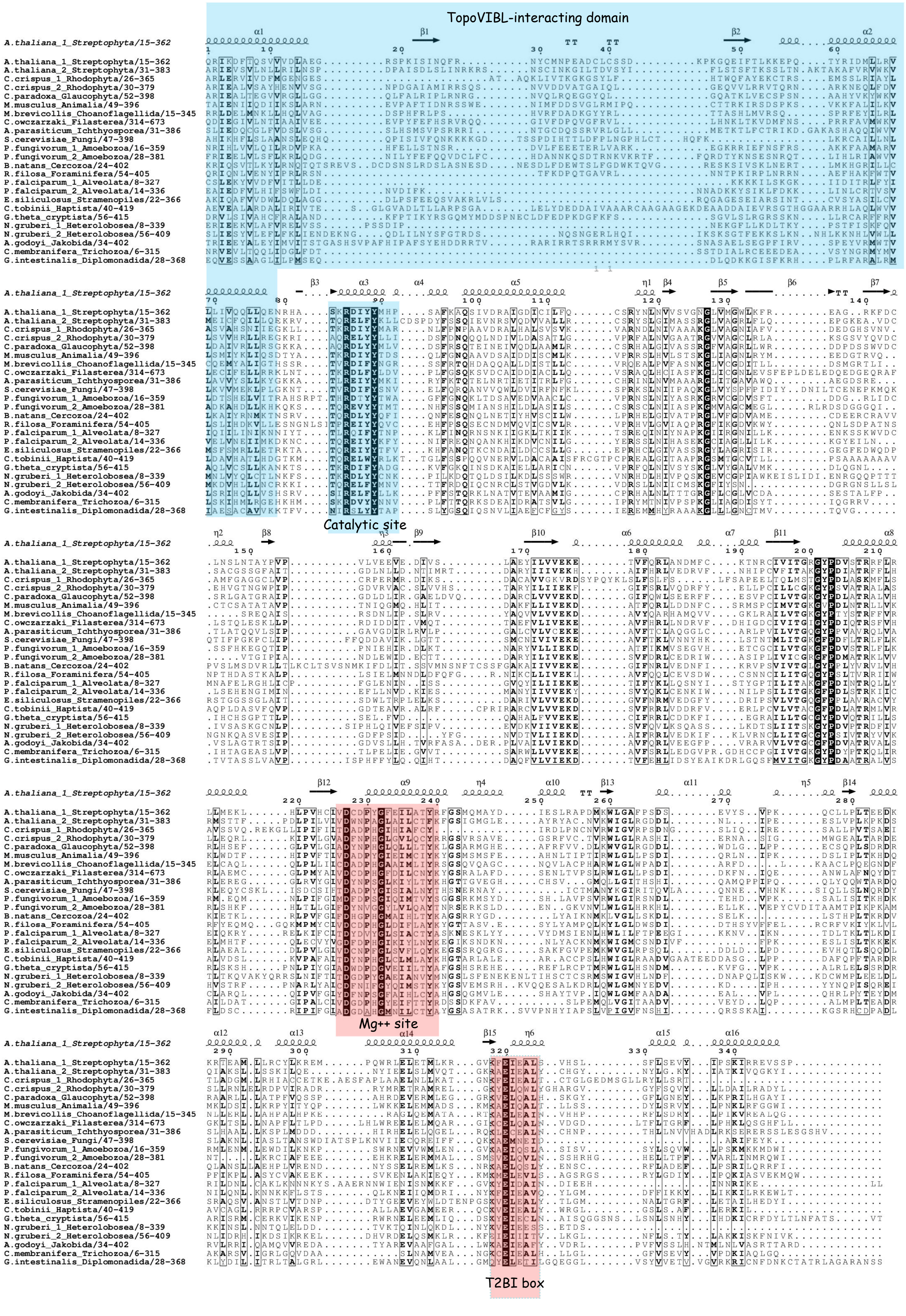
TopoVIBL-interacting domain
Catalytic site
Mg++ site
T2BI box

Figure S8

A

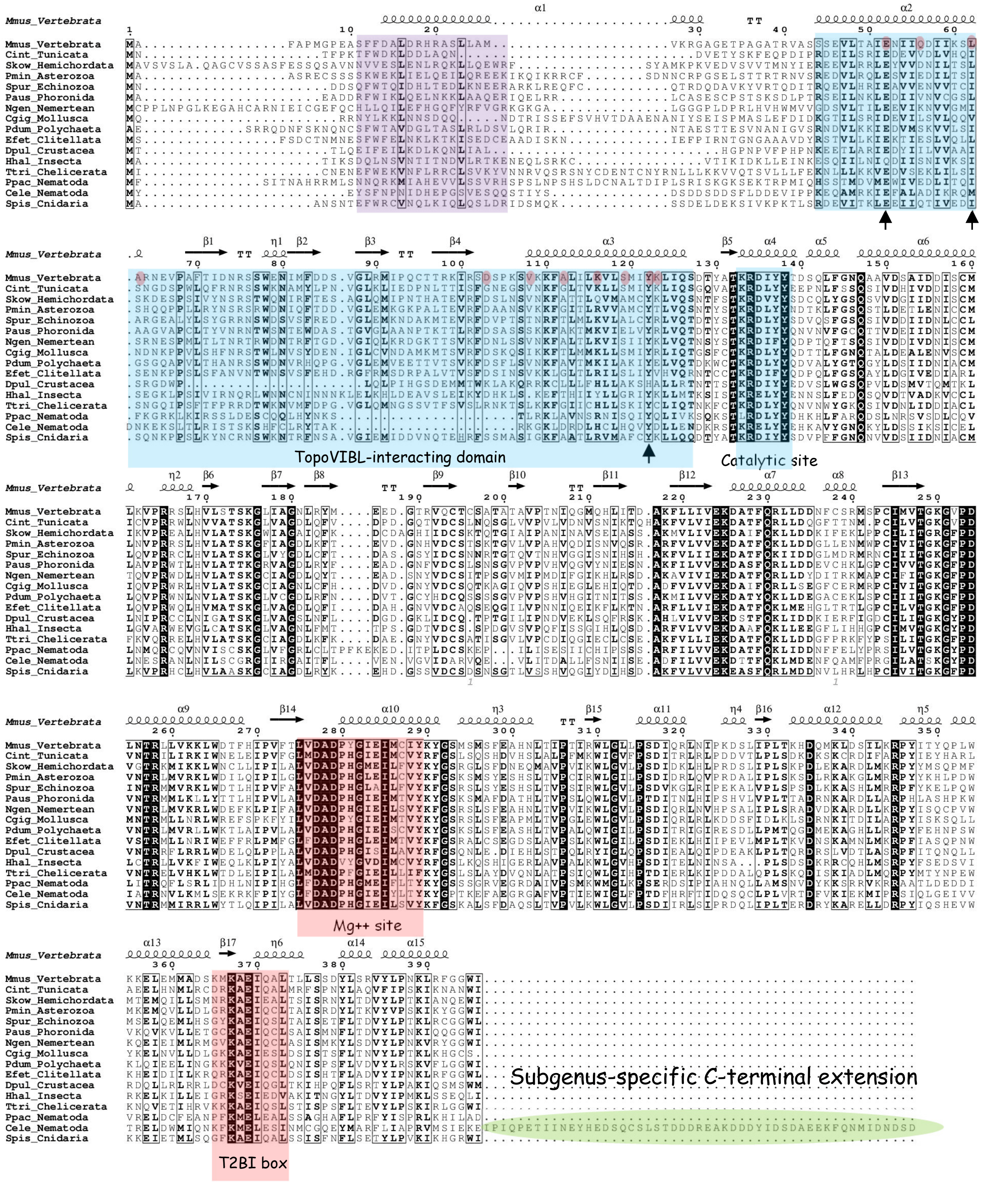
Mmus_Vertebrata
Cint_Tunicata
Skow_Hemichordata
Pmin_Asterozoa
Spur_Echinozoa
Paus_Phoronida
Ngen_Nemertean
Cgig_Mollusca
Pdum_Polychaeta
Efet_Clitellata
Dpul_Crustacea
Hhal_Insecta
Ttri_Chelicerata
Ppac_Nematoda
Cele_Nematoda
Spis_Cnidaria
TopoVIBL-interacting domain
Catalytic site
Mg++ site
Subgenus-specific C-terminal extension
T2BI box

B

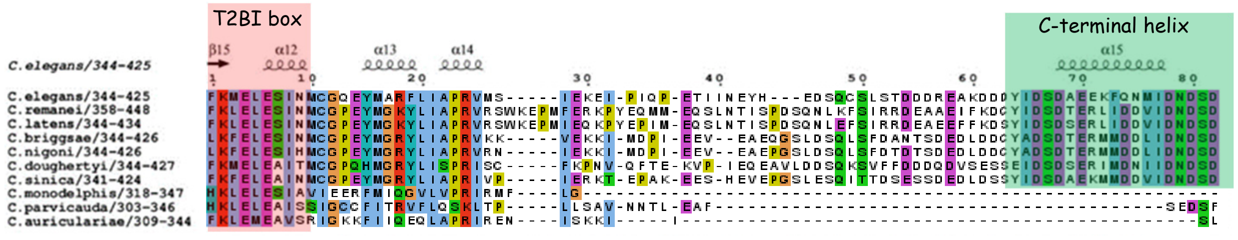
T2BI box
C-terminal helix
C.elegans/344-425
C.remanei/358-448
C.latens/344-434
C.briggsae/344-426
C.nigoni/344-426
C.doughertyi/344-427
C.sinica/341-424
C.monodelphis/318-347
C.parvicauda/303-346
C.auriculariae/309-344

### Figure S9

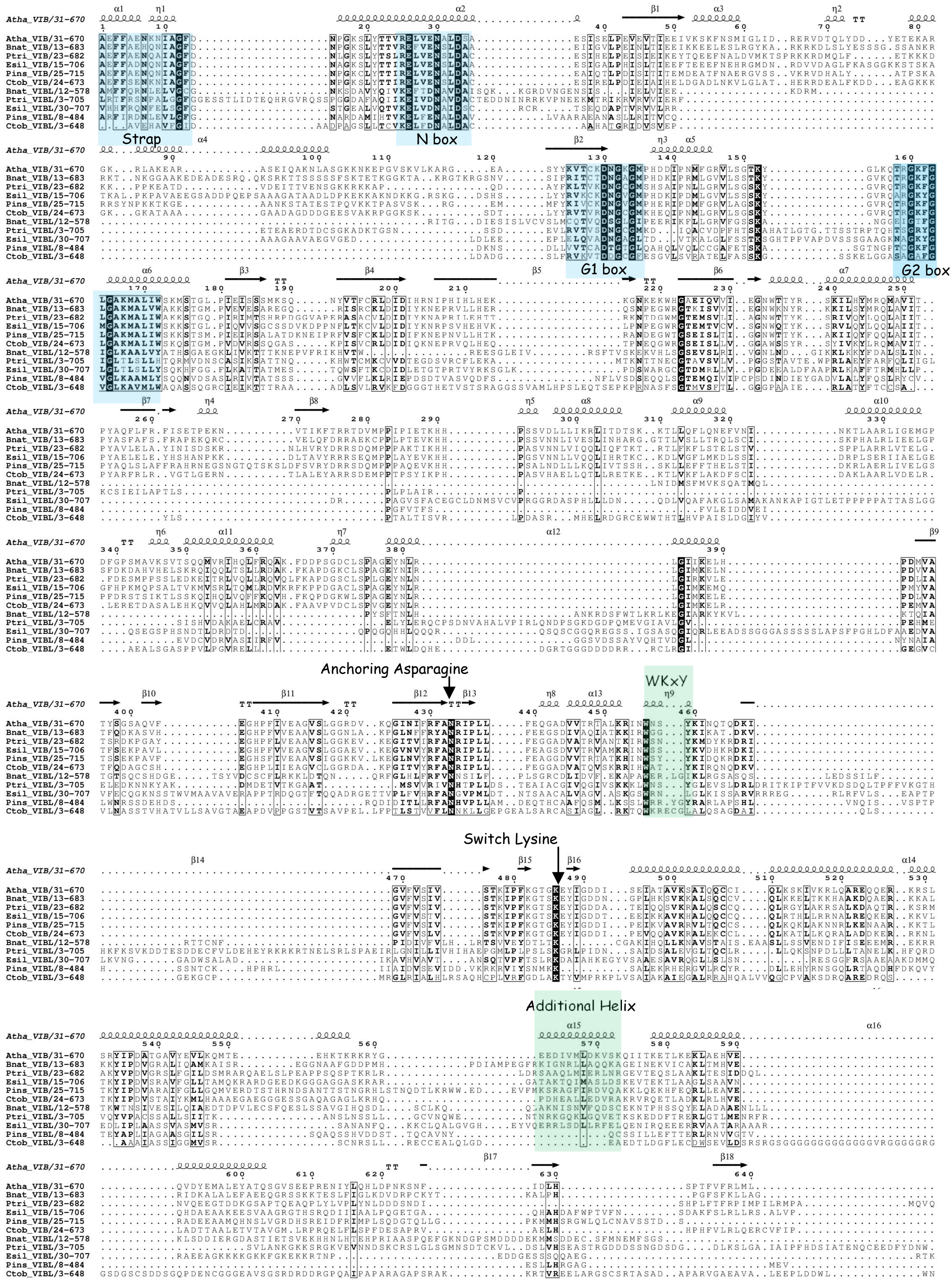
Strap
N box
G1 box
G2 box
Anchoring Asparagine
WKxY
Switch Lysine
Additional Helix

Figure S10

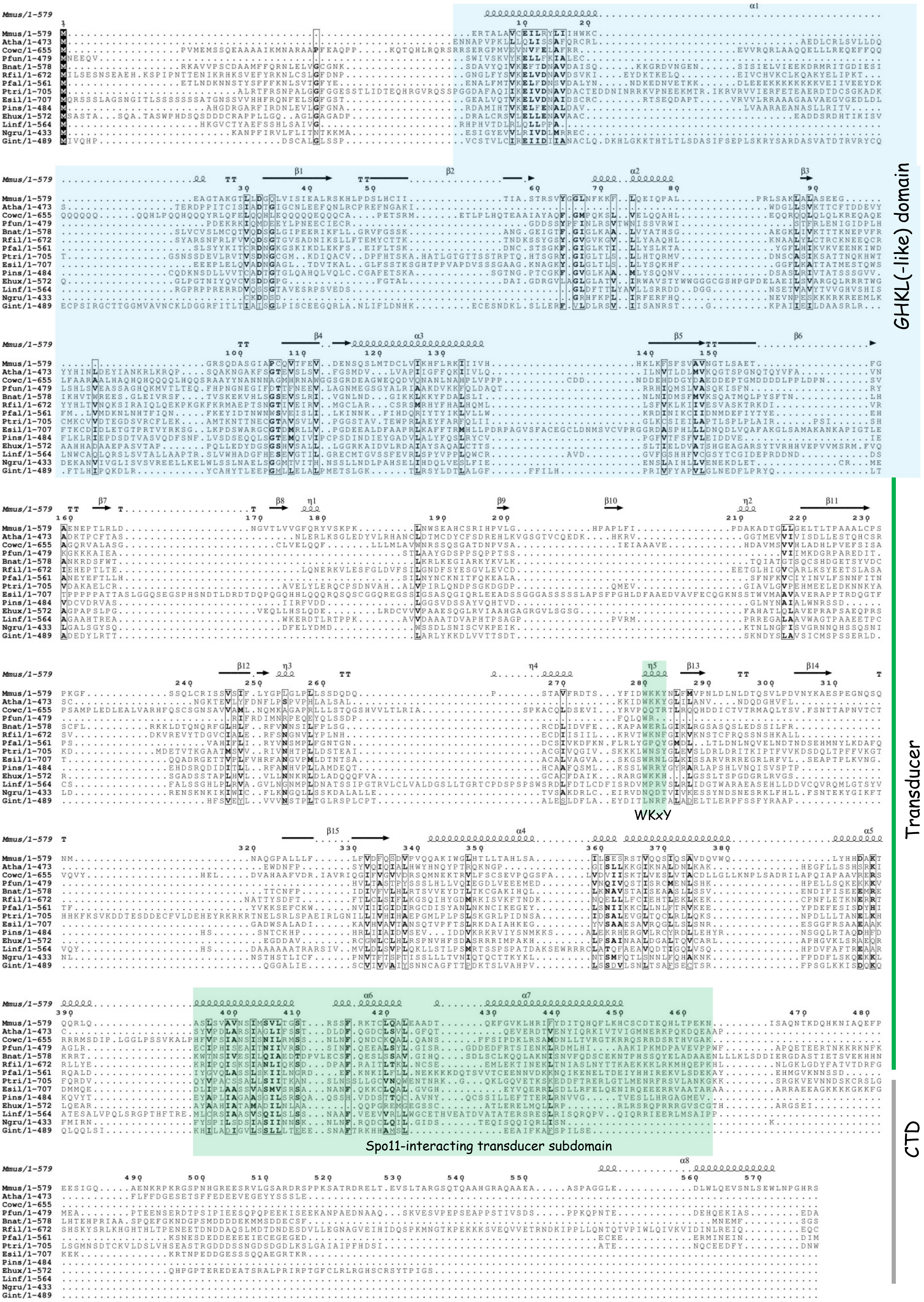

GHKL(-like) domain
Transducer
CTD
WKxY
Spo11-interacting transducer subdomain

### Figure S11

A

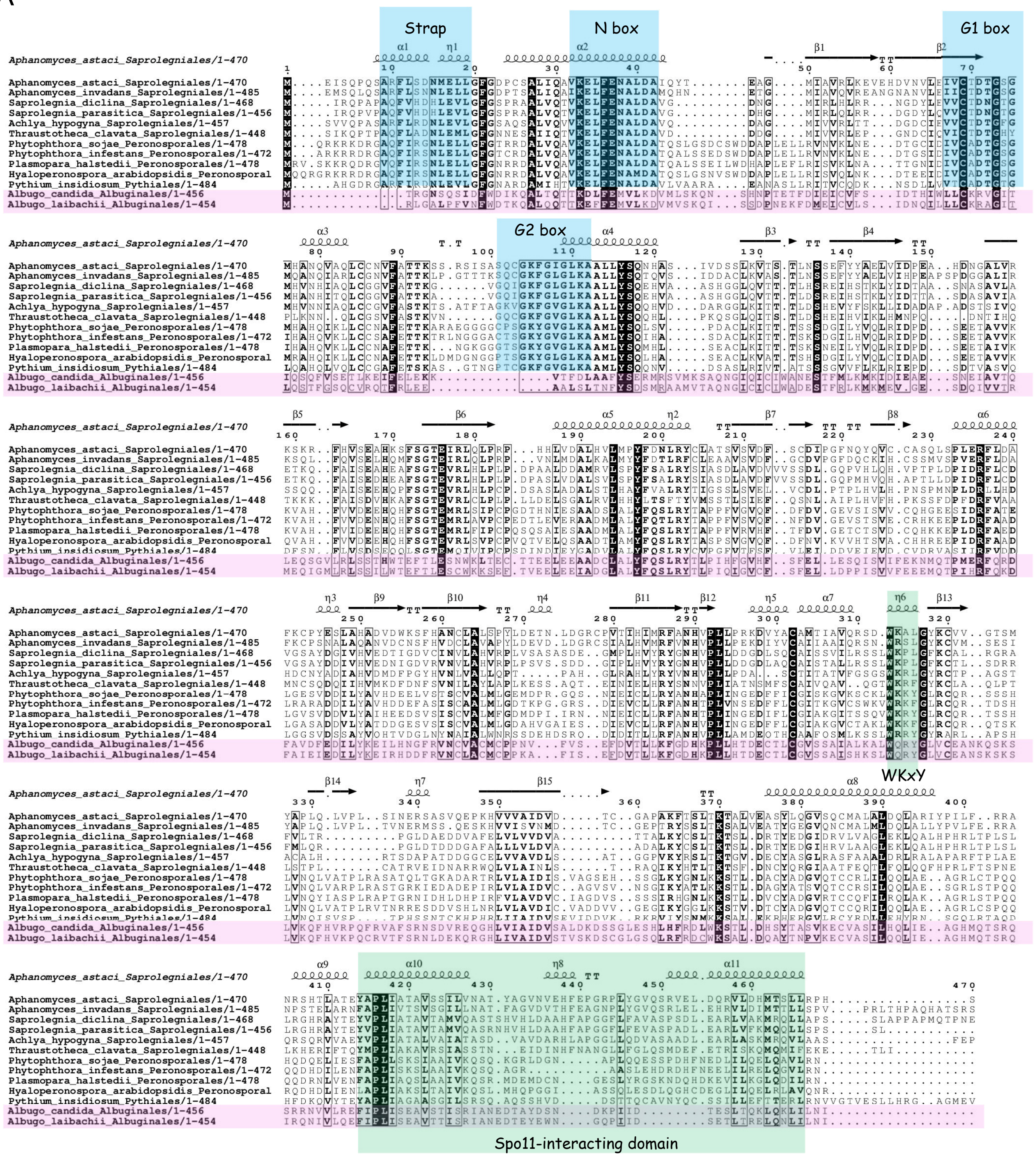


B

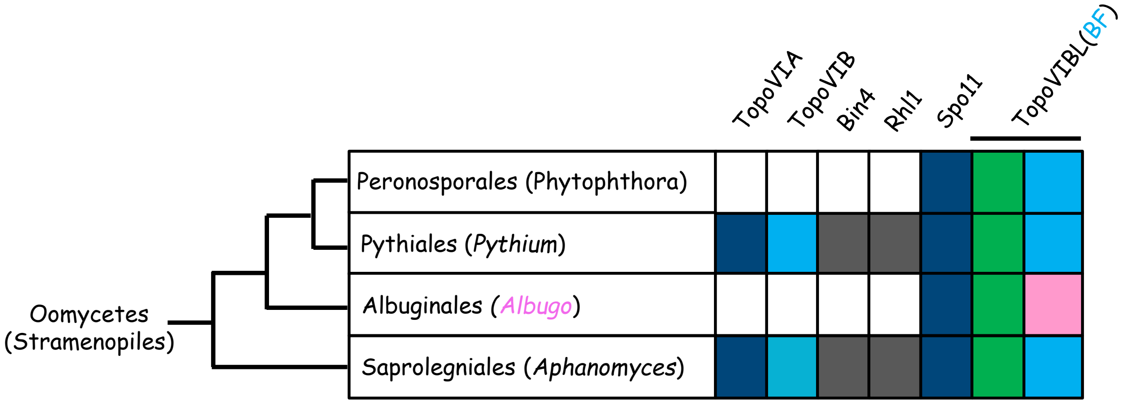

Archaeal vs *Aphanomyces astaci* TopoVIB/VIBL GHKL dimerization

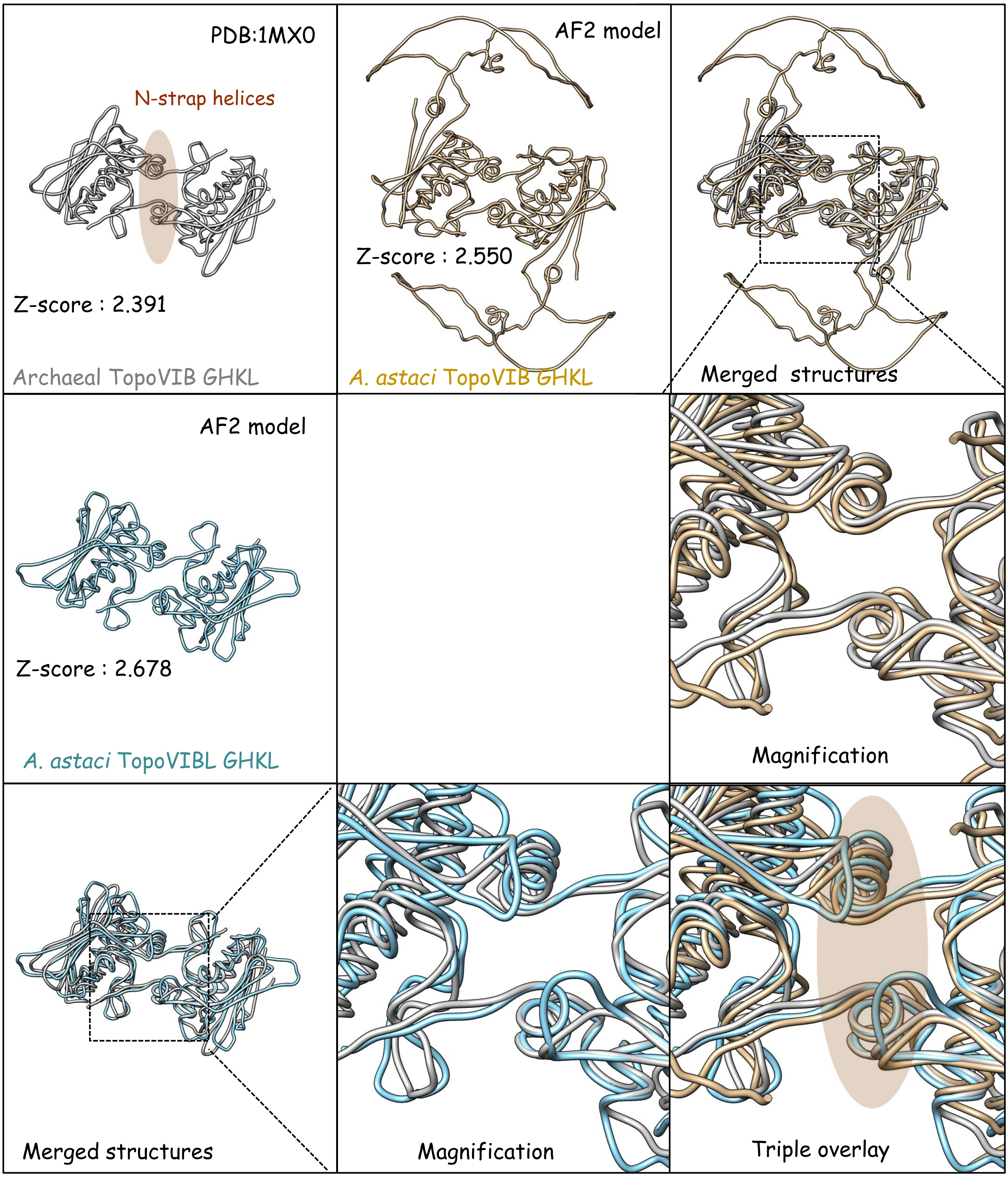

Figure S13

TopoVIB/VIBL-AMP I-TASSER models

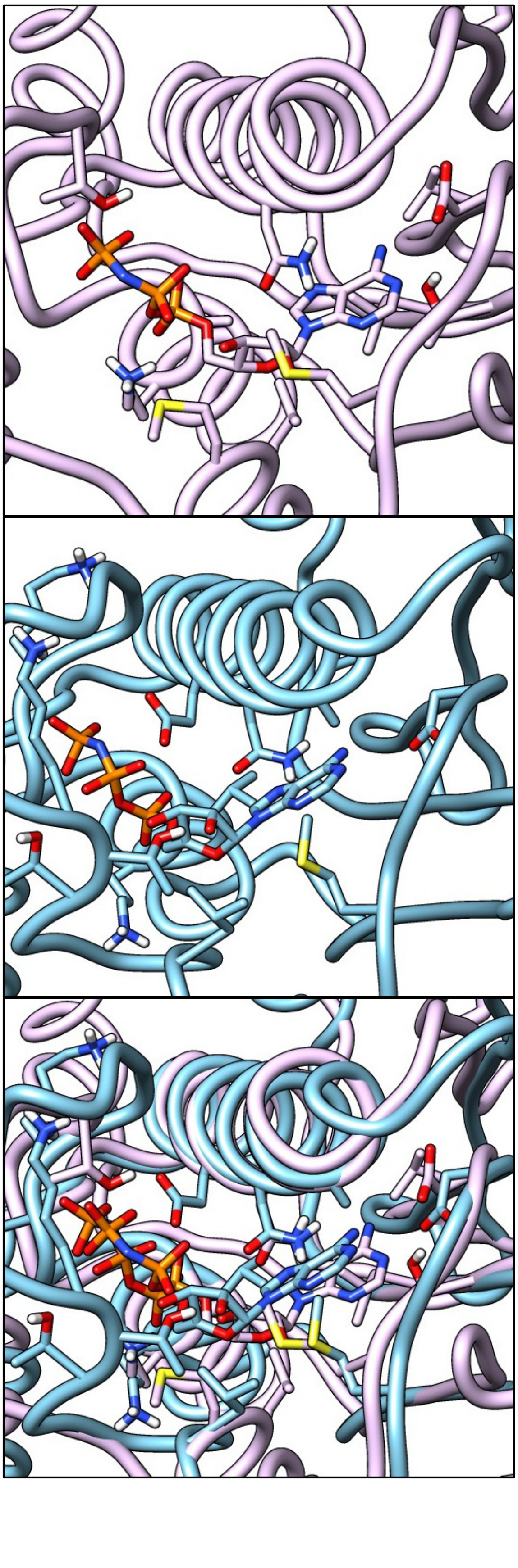

Oomycetal TopoVIB GHKL

Oomycetal TopoVIBL GHKL

Merged structures

Figure S14

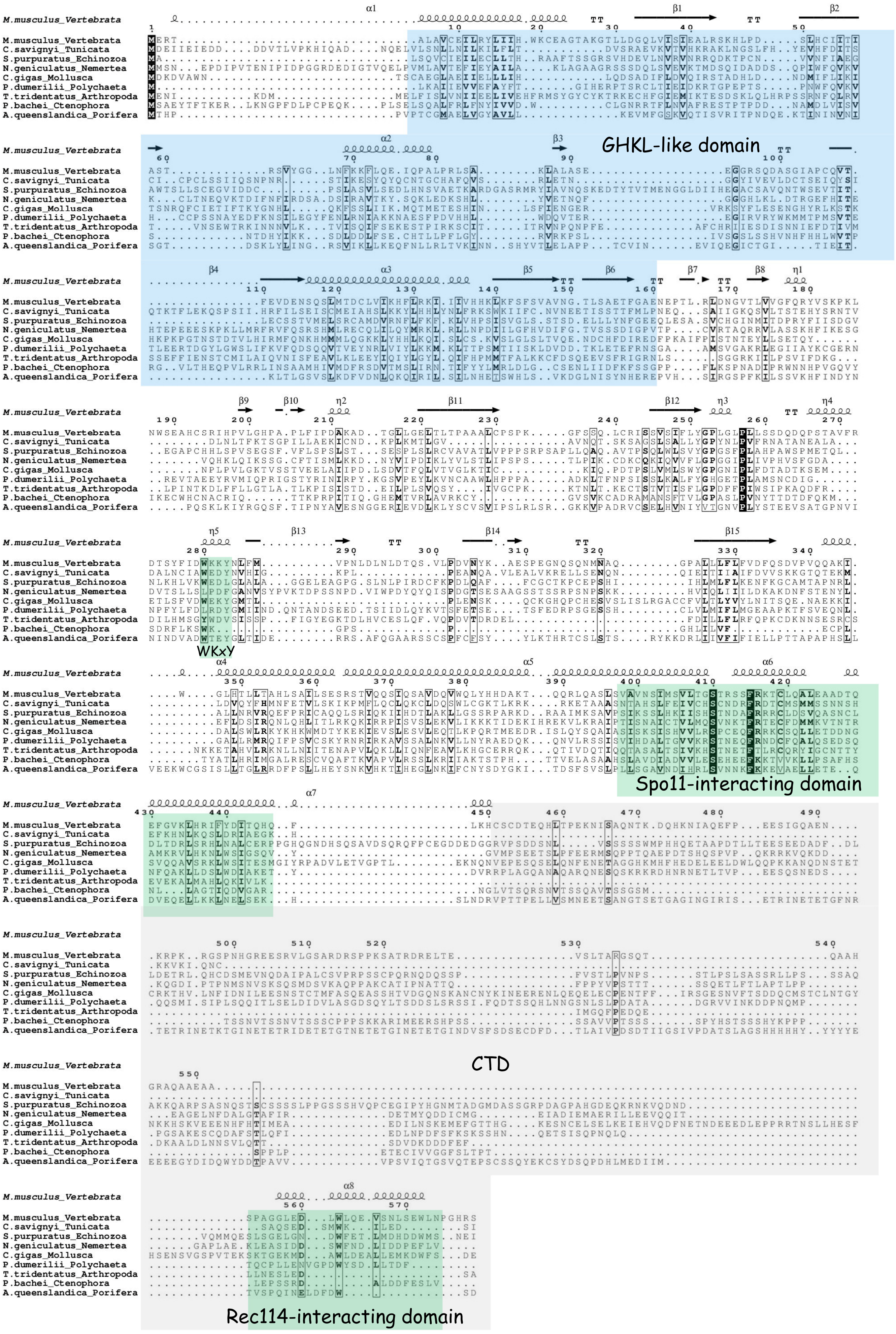
GHKL-like domain
WKxY
Spo11-interacting domain
CTD
Rec114-interacting domain
M.musculus_Vertebrata
C.savignyi_Tunicata
S.purpuratus_Echinozoa
N.geniculatus_Nemertea
C.gigas_Mollusca
P.dumerilii_Polychaeta
T.tridentatus_Arthropoda
P.bachei_Ctenophora
A.queenslandica_Porifera

Figure S15

A

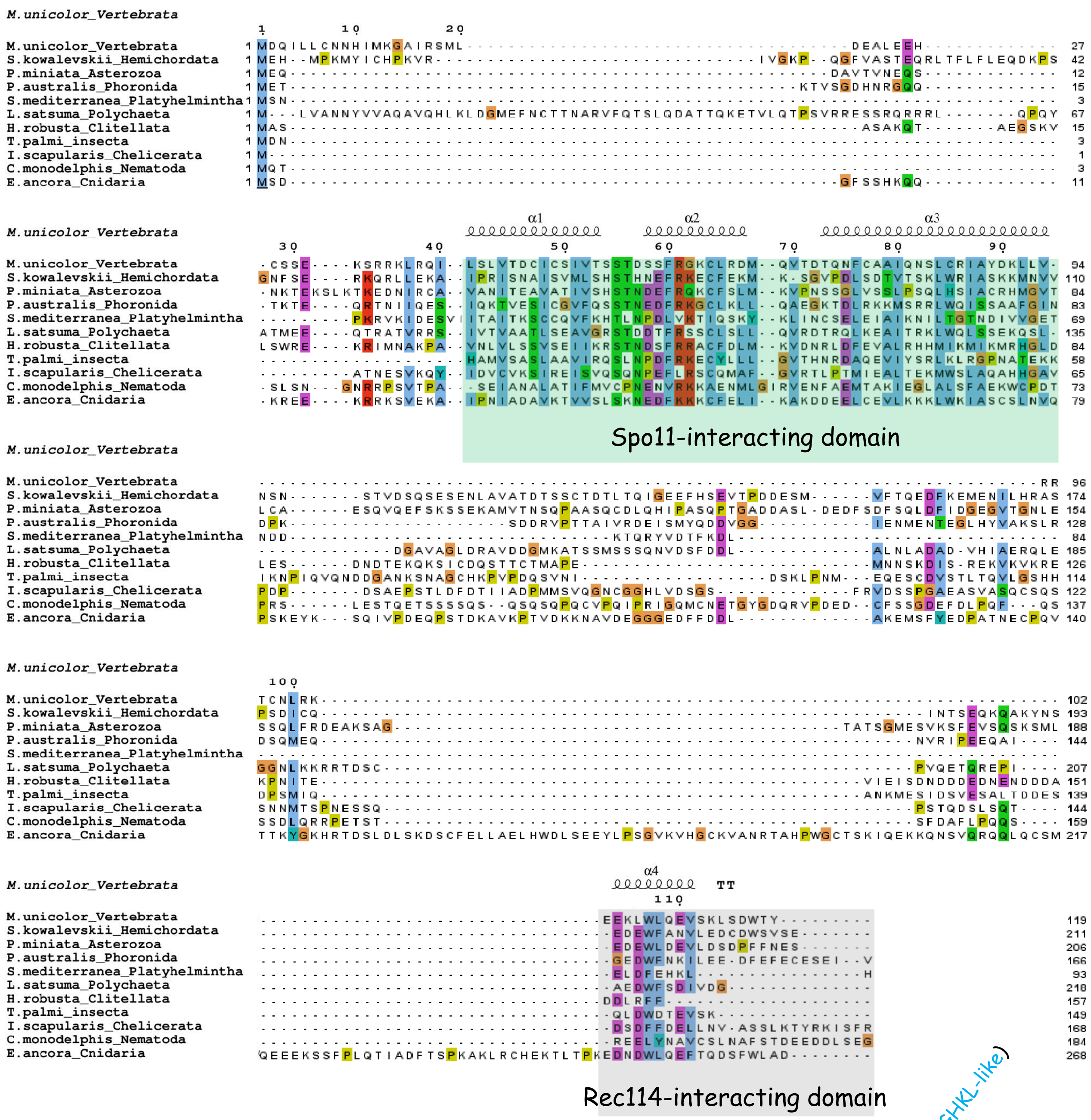


B

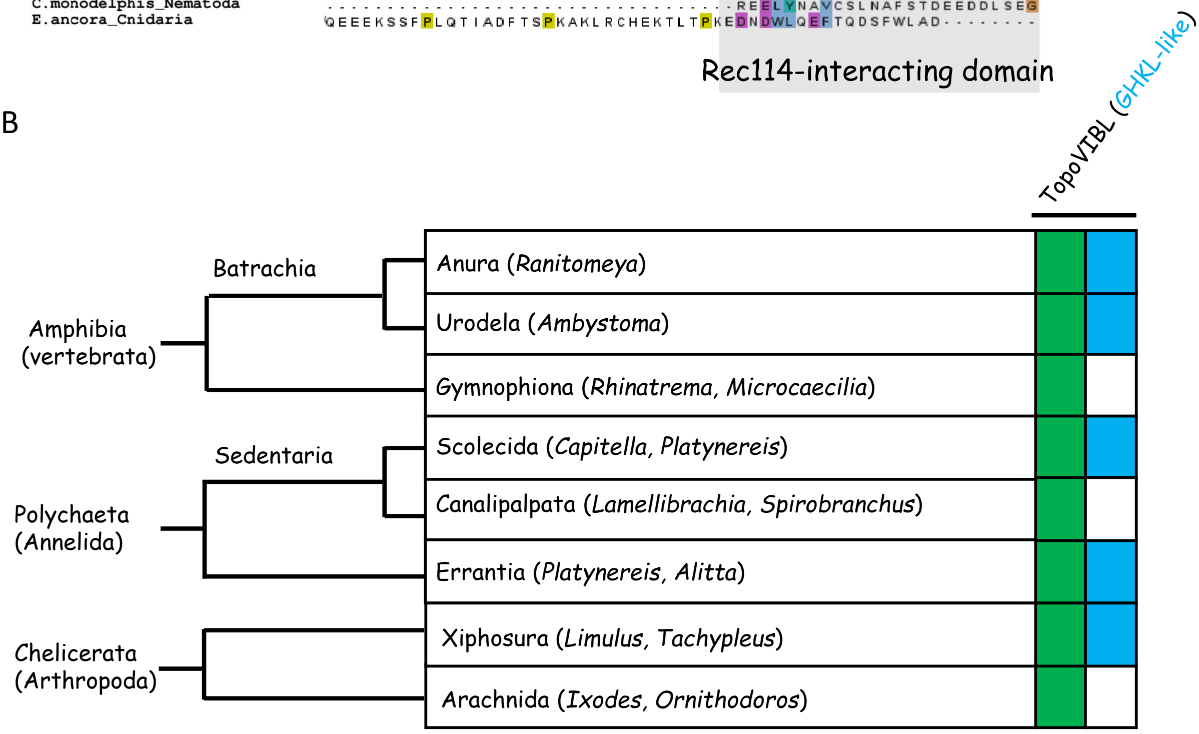

Figure S16

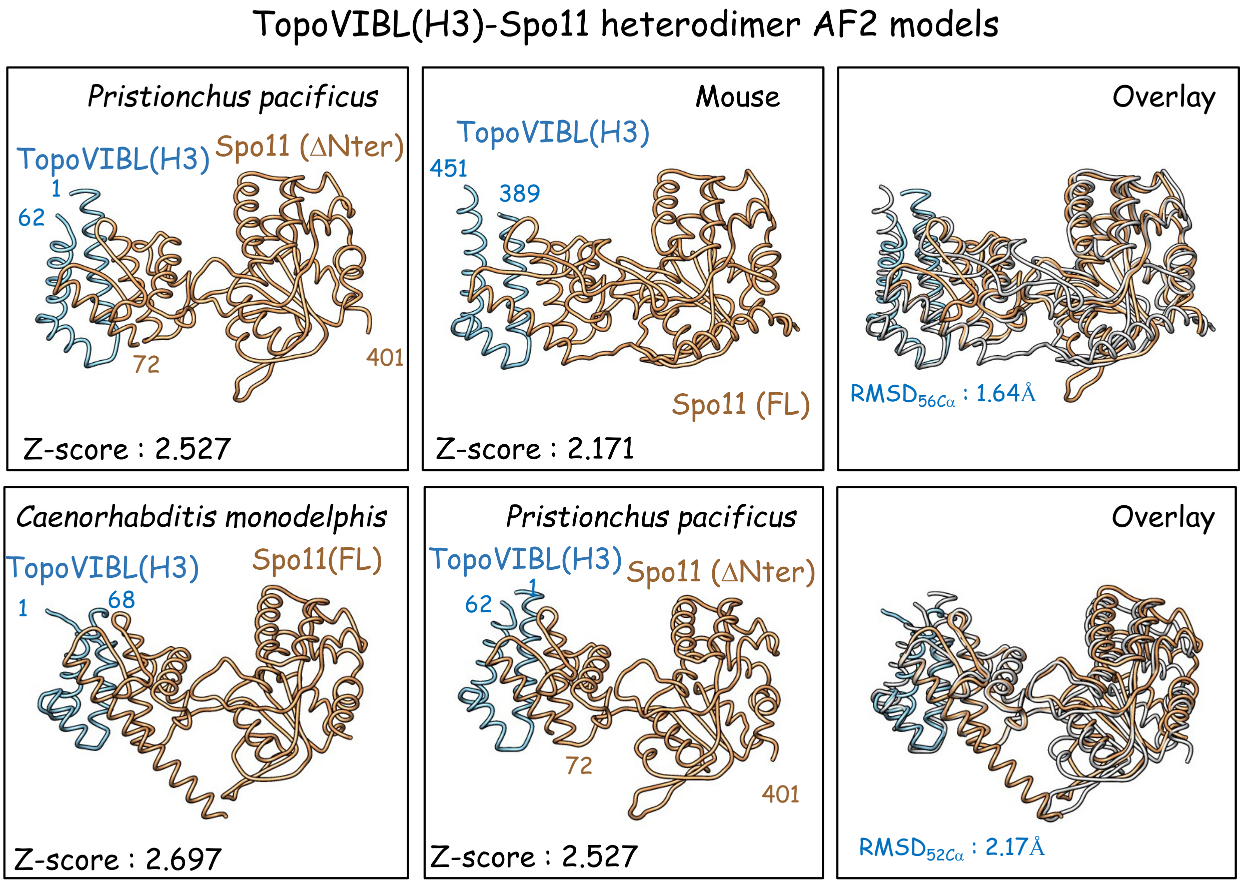
TopoVIBL(H3)-Spo11 heterodimer AF2 models
Pristionchus pacificus
TopoVIBL(H3)
Spo11 (ΔNter)
1
62
72
401
Z-score : 2.527
Mouse
TopoVIBL(H3)
451
389
Spo11 (FL)
Z-score : 2.171
Overlay
RMSD56Cα : 1.64Å
Caenorhabditis monodelphis
TopoVIBL(H3)
Spo11(FL)
1
68
Z-score : 2.697
Pristionchus pacificus
TopoVIBL(H3)
Spo11 (ΔNter)
62
1
72
401
Z-score : 2.527
Overlay
RMSD52Cα : 2.17Å

### Figure S17

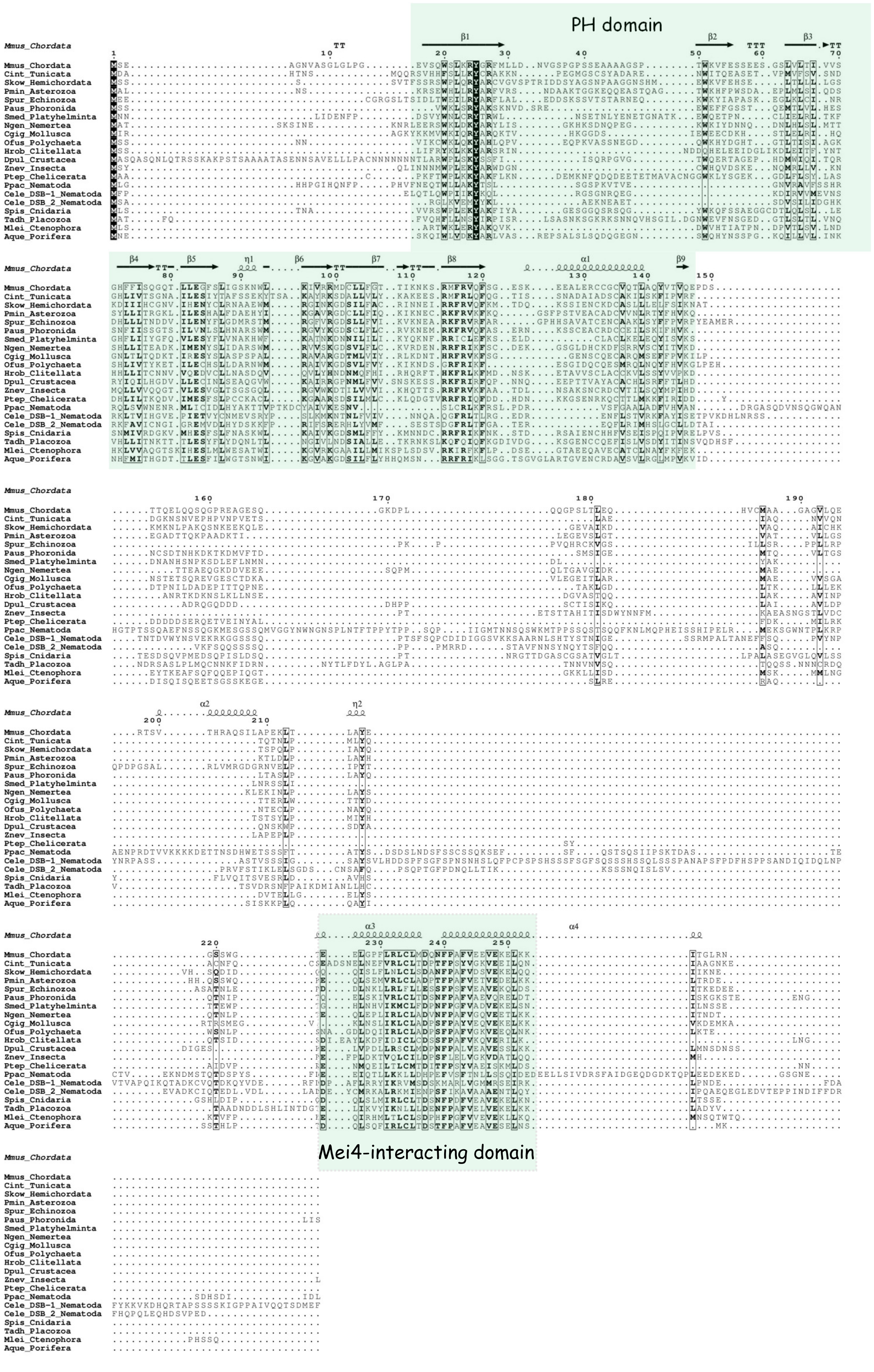

PH domain
Mei4-interacting domain
Mmus_Chordata
Cint_Tunicata
Skow_Hemichordata
Pmin_Asterozoa
Spur_Echinozoa
Paus_Phoronida
Smed_Platyhelminta
Ngen_Nemertea
Cgig_Mollusca
Ofus_Polychaeta
Hrob_Clitellata
Dpul_Crustacea
Znev_Insecta
Ptep_Chelicerata
Ppac_Nematoda
Cele_DSB-1_Nematoda
Cele_DSB_2_Nematoda
Spis_Cnidaria
Tadh_Placozoa
Mlei_Ctenophora
Aque_Porifera

Figure S18

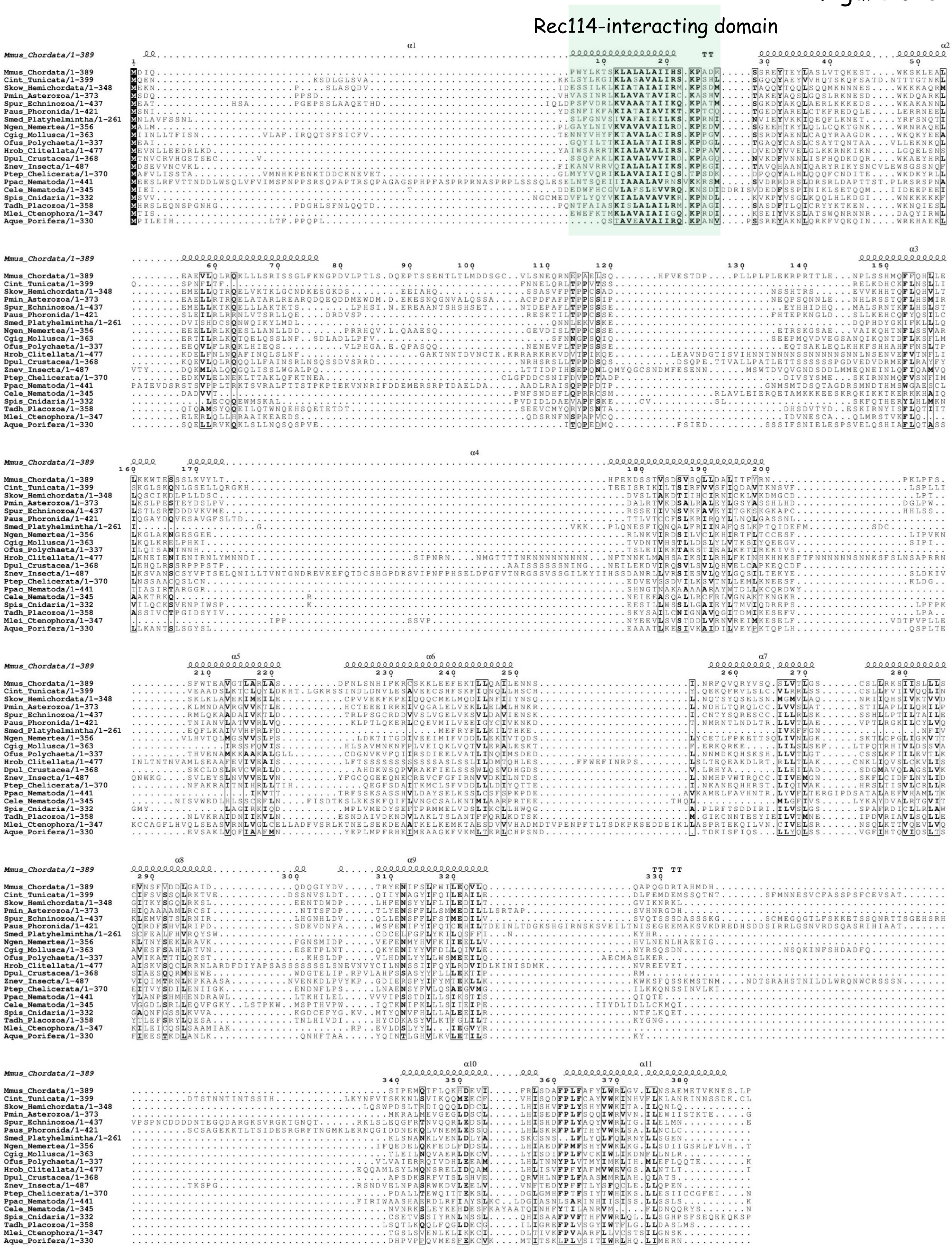
